# Noncanonical Wnt/Ror2 Signaling Regulates Basal Cell Fidelity and Branching Morphogenesis in the Mammary Gland

**DOI:** 10.1101/2025.02.25.640099

**Authors:** Hongjiang Si, Erika Mendoza Mendoza, Madelyn Esquivel, Chad J. Creighton, Jianming Xu, Kevin Roarty

**Affiliations:** Department of Molecular and Cellular Biology, Baylor College of Medicine, Houston, TX 77030; Dan L. Duncan Comprehensive Cancer Center, Breast Cancer Program, Baylor College of Medicine, Houston, TX 77030; Department of Medicine, Baylor College of Medicine, Houston, TX 77030

**Keywords:** Wnt/Ror2 signaling, mammary gland, basal cell identity, branching morphogenesis, epithelial plasticity, RhoA-ROCK1 signaling, cytoskeletal dynamics, lineage transitions, chromatin remodeling, YAP1 signaling

## Abstract

The mammary gland epithelium relies on a delicate balance between basal and luminal cell lineages to maintain tissue homeostasis and enable proper development. While the role of canonical Wnt signaling in mammary biology is well-established, the contribution of noncanonical Wnt signaling to lineage identity has remained unclear. Noncanonical Wnt pathways are primarily associated with morphogenesis, cytoskeletal regulation, and cell migration, but whether they are required for maintaining epithelial cell fate remains largely unexplored. Here, we demonstrate that the noncanonical Wnt receptor Ror2 is expressed in both basal and luminal lineages, yet selectively maintained in basal cells throughout development, suggesting a lineage-specific function. Using a p63^CreERT2/+^ lineage-specific mouse model, we show that Ror2 deletion in basal epithelial cells enhances secondary and tertiary branching while driving a basal-to-luminal fate transition, marked by downregulation of basal markers (K14, K5) and upregulation of luminal markers (K8, K18, ERα). Mechanistically, Ror2 loss disrupts RhoA-ROCK1-YAP1 signaling, leading to cytoskeletal reorganization, chromatin remodeling, and increased accessibility at luminal regulatory loci. Notably, ROCK1 inhibition phenocopies Ror2 loss, reinforcing the critical role of the RhoA-ROCK1 axis in basal cell maintenance. These findings provide direct genetic and mechanistic evidence that noncanonical Wnt signaling is essential for maintaining basal lineage fidelity, offering new insights into the mechanisms regulating epithelial plasticity. Given the fundamental importance of lineage stability in epithelial homeostasis, our results suggest that disruptions in Wnt/Ror2 signaling may contribute to aberrant fate transitions relevant to breast cancer progression.

## Introduction

Epithelial homeostasis is fundamental to maintaining tissue integrity and function, as it ensures a balanced state of cell proliferation, differentiation, and apoptosis, which is crucial for the proper functioning of various organ systems and the prevention of diseases such as cancer. The development of the mouse mammary gland is characterized by remarkable proliferative and differentiation capabilities. This is manifested in the complex physiological processes of ductal extension and branching morphogenesis during pubertal development, the transient remodeling of the gland during estrus cycling, and the proliferation and differentiation of milk-producing alveolar structures during pregnancy^1–3^. The mammary epithelium is primarily composed of two distinct cell types: luminal and basal/myoepithelial. Together, luminal and basal cells of the mammary gland constitute a bi-layered, tubular architecture, a configuration that exhibits its functional efficacy during lactation, wherein the contraction of the outer myoepithelial cells aids in the expulsion of milk from the centrally located alveolar luminal cells^4–6^. Mammary stem cells (MaSCs), positioned at the top of a hierarchical cellular network, play a crucial role in the development of the mammary gland. They possess the unique capability to differentiate into distinct basal and luminal epithelial lineages^7–9^. When transplanted, MaSCs can regenerate a fully functional bilayered ductal tree *in vivo*^7,8^. Lineage-tracing studies have enabled precise labeling of distinct basal and luminal cell lineages, where both basal and luminal layers undergo expansion and self-maintenance via a dedicated pool of unipotent stem cells during puberty and throughout adulthood^10,11^. While there has been considerable progress in understanding the function and regulation of MaSCs, knowledge about the factors that regulate the maintenance of specific lineage cell states remains incomplete^12^. Thus, molecular mechanisms that preserve the identity of luminal or basal epithelial cell states are yet to be fully elucidated.

Wnt/β-catenin signaling orchestrates stem cell renewal and cellular differentiation in various tissues, notably in the mammary gland ^13,14^. This pathway is indispensable from the embryonic stages of mammary development, where it influences mammary placode specification and initiates mammary morphogenesis ^15^, and extends its crucial role to the maintenance of the basal compartment in the adult mammary gland^16^. The developmental trajectory of Wnt/β-catenin responsive MaSCs is intricately modulated by the specific stage of mammary gland development, aligning with the physiological demands of the epithelial network^14^. Distinct from the Wnt/β-catenin pathway, alternative signaling routes exist within the Wnt framework. These noncanonical routes employ unique combinations of Wnt receptors and rely on complex intracellular networks that function independently of β-catenin stabilization^17^. In these noncanonical pathways, key proteins such as small Rho GTPases, JNK, AP-1, and calcium-sensitive enzymes play crucial roles^18^. Although these proteins are not exclusive to Wnt signaling, they are vital for the specialized functions of these alternative pathways. While the role of noncanonical Wnt signaling in guiding polarized cell movement and tissue morphogenesis is well documented, the precise mechanisms through which these diverse Wnt signals influence cell fate decisions and orchestrate the cellular processes involved in morphogenesis remain poorly understood in several organs, including the mammary gland.

Ror2, a receptor tyrosine kinase belonging to the Ror family of Receptor Tyrosine Kinases, harbors an extracellular cysteine-rich domain reminiscent of Frizzled receptors^19^. Ror2-knockout mice exhibit a phenotype akin to Wnt5a-knockout mice, including dwarfism, craniofacial abnormalities, limb defects, and elongated intestines, indicating the involvement of a Wnt5a/Ror2 pathway in various cellular contexts^20^. Our prior research identified the unique expression of Ror2 postnatally within the mammary gland’s epithelial bilayer^21^. We discovered that Ror2 is present in both the luminal and basal epithelial layers, revealing a notable expression gradient: it is pronounced in the differentiated cells of these layers, yet it diminishes in the progenitor and mammary stem cell (MaSC) populations where Wnt/β-catenin signaling predominates. The distinct and sustained expression of Ror2 in mature epithelial cells, particularly within the basal lineage, prompted a detailed investigation into its lineage-specific roles *in vivo*.

In this study, we aimed to investigate the role of Ror2 in regulating basal epithelial cell homeostasis *in situ*, noting its consistent expression throughout this cellular compartment across postnatal developmental stages^21^. Utilizing a novel basal lineage-specific p63^CreERT2/+^ knock-in mouse model in combination with a Ror2^f/f^ conditional mutant mouse, we achieved lineage-specific and temporal deletion of Ror2 *in vivo*. Unexpectedly, the conditional deletion of Ror2 in basal cells led to an increase in branching morphogenesis and a basal-to-luminal cell fate switch *in situ*. The descendants of Ror2-deleted basal cells predominantly became ER^+^ luminal cells, as opposed to ER-negative luminal cells, indicating that Ror2 status influences lineage specification. Single-cell ATAC-sequencing revealed that upon Ror2 loss, basal cells segregated into three distinct clusters with luminal and basal chromatin characteristics, encompassing basal, luminal progenitor, and mature luminal subsets. In contrast, wild-type basal cells retained a basal-oriented gene program within a single cluster. These novel insights underscore the critical role of Wnt/Ror2 signaling in basal cell lineage maintenance and highlight the broader impact of our findings on understanding epithelial cell differentiation and homeostasis.

## Results

### Basal lineage-specific Ror2 signaling controls branching morphogenesis

Given the consistent expression of Ror2 across developmental stages within the basal epithelial layer of the mammary gland^21^, we used a lineage-specific p63^CreERT2/+^ knock-in mouse model in combination with a Ror2^f/f^ conditional mutant mouse and ROSA^mTmG^ two-color fluorescent Cre-reporter mouse to genetically deplete Ror2 *in vivo* (**Figure 1A**). To investigate the functional consequences of Ror2-deficient basal cells on mammary gland development, we administered Tamoxifen (TAM, 80 µg/g body weight) to pubertal postnatal day 42 (P42) control p63^CreERT2/+^;ROSA^mTmG/+^ and experimental p63^CreERT2/+^;ROSA^mTmG/+^; Ror2^fl/fl^ female mice for 2 consecutive days to induce recombination (**Figure 1B**). A 48-hour timepoint was chosen post-TM administration to verify lineage-specific recombination and deletion of Ror2 *in situ* within the basal epithelial layer of the mammary gland. Following administration of TM, successful recombination was confirmed by the expression of membrane eGFP in control p63^CreERT2/+^;ROSA^mTmG/*+*^ mice, indicating basal-specific distribution of eGFP (**Figure 1C**). In experimental p63^CreERT^^2/+^;ROSA^mTmG/+^; Ror2^fl/fl^ mice, eGFP expression was also localized specifically within the basal epithelial layer (**Figure 1D**), accompanied by the targeted deletion of Ror2, as demonstrated at the level of immunofluorescence (IF) and flow cytometry. IF analysis revealed the loss of Ror2 expression in eGFP^+^ basal cells of the experimental mice, confirming effective recombination and gene deletion (**Figure 1E-F**). Additionally, we validated the deletion of Ror2 by performing quantitative PCR (qPCR) on eGFP^+^ cells isolated from the mammary glands of mTmG reporter mice, showing a significant reduction of Ror2 mRNA levels in eGFP^+^ cells from the experimental mice compared to controls (**Figure 1G**). These results collectively confirm the successful recombination and basal-specific deletion of Ror2 *in vivo*.

**Figure 1.**
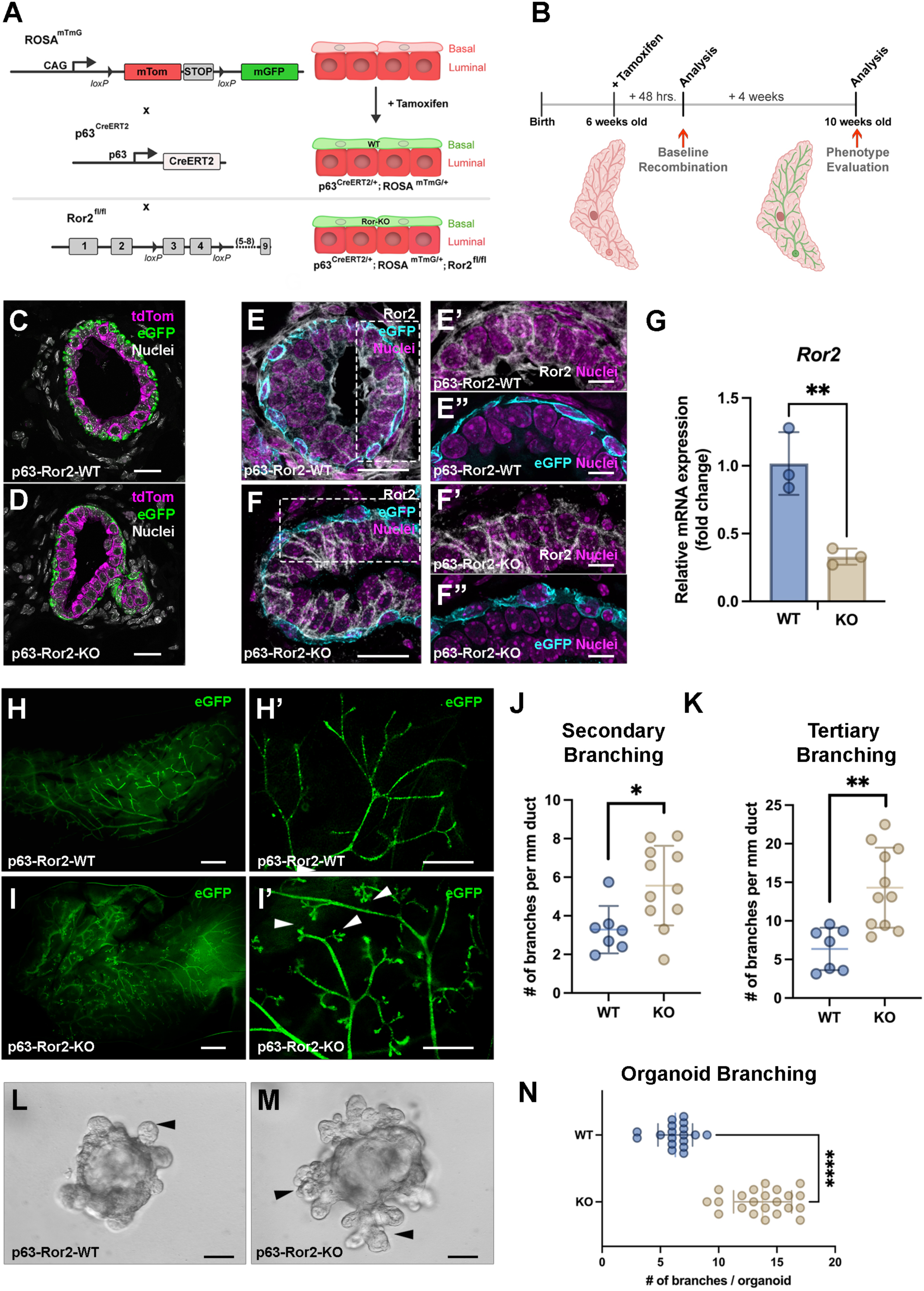
Temporal deletion of Ror2 in p63^+^ basal epithelial cells enhances mammary gland branching morphogenesis *in vivo* and *in vitro*. (A) Schematic illustrating the breeding strategy used to generate tamoxifen-inducible p63-specific Ror2 knockout mice carrying a heterozygous ROSA^mTmG^ reporter allele. (B) Experimental timeline showing tamoxifen administration, tissue collection, and phenotype analysis. (C, D) Representative immunofluorescence images of ROSA^mTmG^-tdTomato (magenta) and ROSA^mTmG^-eGFP (green) in mammary gland cross-sections from (C) control and (D) p63-Ror2-KO mice 2 days post-tamoxifen injection. Scale bar: 20 μm. (E, F) Representative immunofluorescence images showing Ror2 protein (gray) by immunofluorescence in mammary gland cross-sections from (E) control and (F) p63-Ror2-KO mice at 2 days post-tamoxifen injection. Scale bar: 20 μm. (E’-E”) Magnified inset of E showing Ror2 expression within the p63Cre-eGFP^+^ basal layer (cyan) in controls and (F’-F”) Ror2 –KO glands. Scale bar: 10 μm. (G) Quantitative RT-qPCR analysis of Ror2 mRNA expression in FACS-isolated mammary basal epithelial cells from control and p63-Ror2-KO mice. Expression levels were normalized to GAPDH, with data represented as fold change relative to the control group (p = 0.0076; n = 3 biological replicates per group). (H-H’) Representative whole-mount epifluorescence images of eGFP in mammary gland structures from control mice. (H) Wholemount mammary gland (scale bar: 2 mm) and (H’) a magnified view of ducts (scale bar: 1 mm). (I, I’) Representative whole-mount epifluorescence images of eGFP in mammary gland structures from p63-Ror2-KO mice. (I) Whole mammary gland (scale bar: 2 mm) and (I”) a magnified view of ducts (scale bar: 1 mm). Increased branching in p63-Ror2-KO mammary glands is indicated by white arrowheads. (J, K) Quantification of (J) secondary branching and (K) tertiary branching in mammary glands from control and p63-Ror2-KO mice (p = 0.0184 and p = 0.0019, respectively; n = 6 glands for control and n = 11 glands for p63-Ror2-KO groups). (L, M) Representative differential interference contrast (DIC) bright-field images of branching mammary organoids from control and p63-Ror2-KO groups at 72 hours post-seeding with FGF stimulation. Increased branching morphogenesis in p63-Ror2-KO organoids is indicated by black arrowheads. Scale bar: 50 μm. (N) Quantification of the number of branching projections per organoid in control and p63-Ror2-KO groups (p < 0.0001; n = 18 organoids for control and n = 21 organoids for p63-Ror2-KO).

We proceeded to investigate the consequences of p63-specific Ror2 deletion on ductal morphogenesis. Cre recombination was initiated during puberty at P42 (6 weeks) of age in both control p63^CreERT2/+^;ROSA^mTmG/+^ and experimental p63^CreERT2/+^;ROSA^mTmG/+^; ^Ror2fl/fl^ mice through the administration of TM, to assess the impact of Ror2 loss on ductal extension and branching during this active phase of ductal development. To evaluate the impact of Ror2 deletion, the #3 and #4 inguinal mammary glands were examined 4 weeks post-recombination, at postnatal day 70 (10 weeks). We observed that p63^CreERT2/+^;ROSA^mTmG/+^; Ror2^fl/fl^ glands exhibited an equivalent amount of ductal extension compared to control p63^CreERT2/+^;ROSA^mTmG/+^ mammary glands (**Figure 1H, I**). Although primary branch points were consistent between control Ror2-intact glands and experimental Ror2-deficient glands, a statistically significant increase in secondary (**Figure 1J**) and higher order tertiary branching (**Figure 1K**) occurred in Ror2-deficient mammary glands compared to controls. In addition to *in vivo* analyses, we examined branching morphogenesis using an organotypic culture model derived from p63^CreERT2/+^;ROSA^mTmG/+^ and p63^CreERT2/+^;ROSA^mTmG/+^; Ror2^fl/fl^ mammary glands following a 2-day administration of TM *in vivo* followed by the isolation of recombined epithelial fragments from the glands. In this model, epithelial cells were isolated, lineage-depleted, and then embedded in growth factor-reduced Matrigel. Following the *in vitro* propagation of organotypic cultures after timed addition of FGF2 over 1 week, an increase in branching was evident in Ror2-deficient organoids compared to controls, consistent with *in vivo* findings (**Figure 1L-M**). Collectively, these results demonstrate that the basal-specific loss of Ror2 signaling significantly enhances branching morphogenesis of the mammary gland during ductal extension and patterning from puberty into adulthood.

### Wnt/Ror2 signaling regulates basal epithelial identity *in vivo*

To better assess the phenotypic differences in morphology between p63^CreERT2/+^;ROSA^mTmG/+^ and p63^CreERT2/+^;ROSA^mTmG/+^;Ror2^fl/fl^ mammary glands, we performed wholemount confocal imaging of adult mammary glands that were rendered optically clear by CUBIC-based approaches^22,23^. We utilized the double-fluorescent Cre reporter mouse model and membrane-targeted GFP (mG) post-excision to achieve precise visualization of the ductal architecture and discern the cellular morphologies of both control and Ror2-deficient basal cells within the mammary gland. In control p63^CreERT2/+^;ROSA^mTmG/+^ mammary glands, elongated spindle shaped GFP-positive basal cells could be visualized wrapping around the epithelial ducts, positioned adjacent to the mammary stroma (**Figure 2A, A’**). Notably, in Ror2-deficient mammary glands, GFP-positive basal cells exhibited a diminished spindle-shaped morphology along with discontinuous coverage of the ducts, compared to control basal cells, highlighting the morphological impact of Ror2 deletion. (**Figure 2B, B’**)

**Figure 2.**
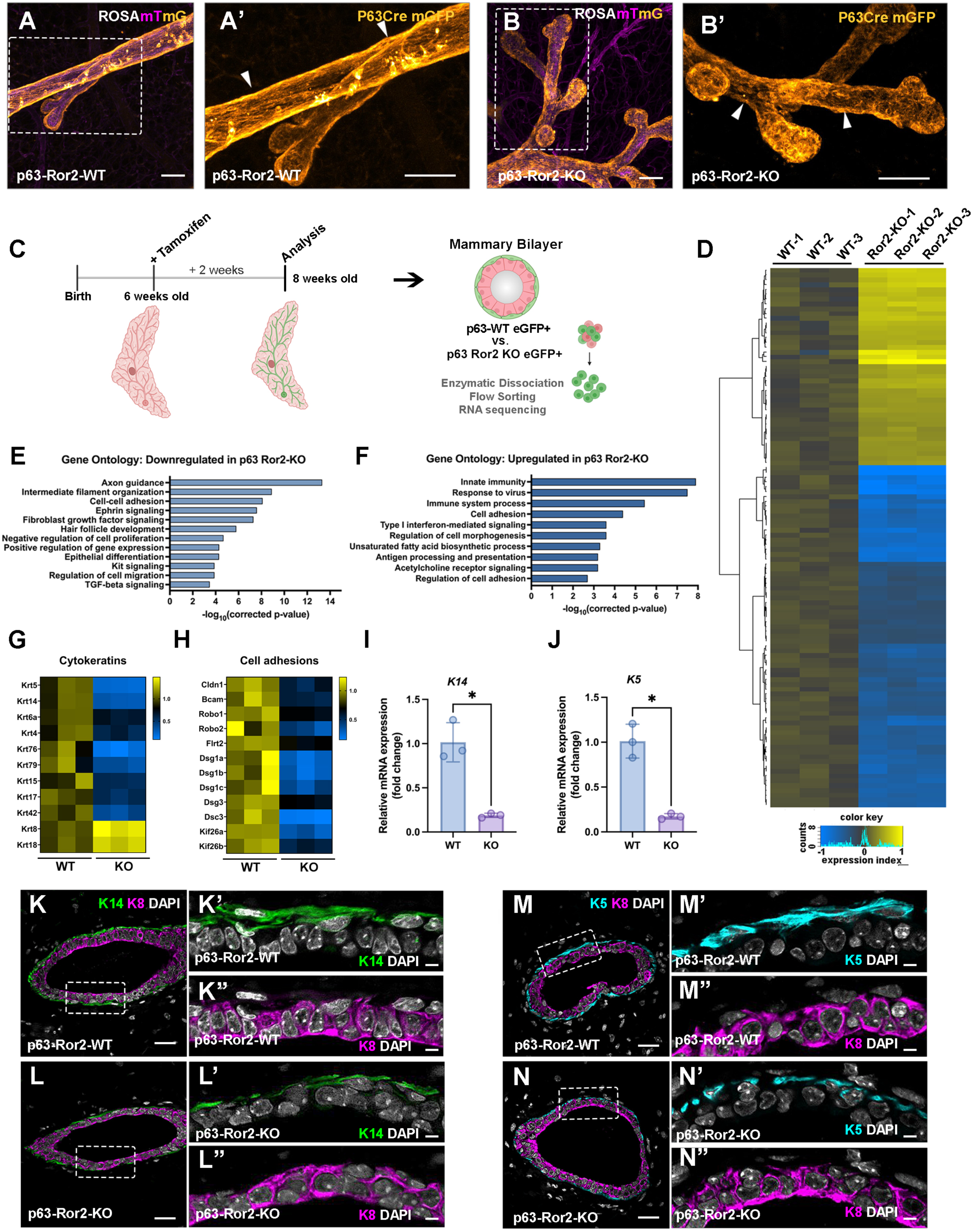
Ror2 loss in basal cells alters cytokeratin expression and adhesion gene expression programs. (A, B) Confocal Maximum Intensity Projection (MIP) of mammary ducts from ROSA^mTmG^ glands from (A, A’) p63-Cre WT and (B, B’) p63-Cre-Ror2-KO glands optically cleared with CUBIC. Scale bars: 70 μm. (magenta, unrecombined mTomato; orange, recombined mGFP). (C) Schematic of experimental timeline showing tamoxifen administration and tissue collection for RNA sequencing. (D) Heatmap of significantly differentially expressed genes (adjusted p < 0.05 and fold change > 1.5) in control and p63-Ror2-KO basal cells, as identified by RNA sequencing. Expression levels are presented as log_2_(reads/average reads in the control group) and visualized using a gradient from blue (downregulation) to yellow (upregulation). (E, F) Gene ontology analysis performed using the DAVID Bioinformatics Database, highlighting the enrichment of gene expression changes (E) downregulated and (F) upregulated in several biological processes following Ror2 loss. (G, H) Heatmaps displaying RNA sequencing results for differentially expressed genes related to (G) cytokeratin and (H) cell adhesion. Expression changes are represented as fold changes using a gradient from blue (downregulation) to yellow (upregulation). (I) Quantitative RT-qPCR analysis of *K14* mRNA expression in FACS-isolated basal epithelial cells from control and p63-Ror2-KO mice. Gene expression levels were normalized to *GAPDH*, and data are presented as fold change relative to the control group (p = 0.0030; n = 3 biological replicates per group). (J) Quantitative RT-qPCR analysis of *K5* mRNA expression in FACS-isolated basal epithelial cells from control and p63-Ror2-KO mice. Gene expression levels were normalized to *GAPDH*, and data are presented as fold change relative to the control group (p = 0.0016; n = 3 biological replicates per group). (K, L) Representative immunofluorescence images of K14 (green) and K8 (magenta) in mammary gland cross-sections from (K) control and (L) p63-Ror2-KO mice at 4 weeks post-tamoxifen injection. Scale bar: 25 μm. (K’, K”, L’, L”) Magnified inset of single-channel images of K14 and K8 reveal reduced K14 coverage in p63-Ror2-KO mammary ducts. Scale bar 10 μm. (M, N) Representative immunofluorescence images of K5 (cyan) and K8 (magenta) in mammary gland cross-sections from (M) control and (N) p63-Ror2-KO mice at 4 weeks post-tamoxifen injection. Scale bar: 25 μm. (M’, M”, N’, N”) Magnified single-channel images of K5 and K8 reveal reduced K5 coverage in p63-Ror2-KO mammary ducts relative to p63 WT ducts.

To determine the molecular underpinnings of Ror2-mediated effects in the basal mammary epithelial layer, we performed bulk RNA sequencing of sorted GFP-positive cells from p63^CreERT2/+^;ROSA^mTmG/+^ and p63^CreERT2/+^;ROSA^mTmG/+^;Ror2^fl/fl^ mammary glands at 8 weeks (about 2 months) of age following a 2-week chase post TM-mediated Cre recombination in control and experimental glands (**Figure 2C**). The deletion of Ror2 within basal cells resulted in the upregulation of 328 genes and the downregulation of 297 genes (**Figure 2D**). Interestingly, when we applied gene ontology analysis to the list of altered genes, the top represented terms among downregulated genes included those associated with axon guidance, cell adhesion, intermediate filament organization, and response to mechanical stimulus (**Figure 2E**). Genes upregulated were represented by categories associated with cell morphogenesis and the immune system (**Figure 2F**). The critical role of Wnt/Ror2 signaling in maintaining basal-specific functions within the mammary gland was illustrated by the marked downregulation of basal-specific cytokeratins, including Keratin 5, Keratin 14, and Keratin 17, alongside desmosomal adhesion proteins such as *Dsg1a-c*, *Dsg3*, and *Dsc3* – key mediators of barrier function and mechanical integrity in the myoepithelial layer (**Figure 2G-H, S1**). This downregulation of basal-specific cytokeratins was confirmed both at the histological level by immunofluorescence and at the transcriptional level by qPCR (**Figure 2I-N**). In contrast, the striking upregulation of luminal-specific cytokeratins K8 and K18 suggested a potential shift in cell identity (**Figure 2G**), prompting further investigation into cell state changes downstream of Wnt/Ror2 loss-of-function. Of note, *Kif26b*, a kinesin family protein previously identified as a Ror2 target in Mouse Embryonic Fibroblasts (MEFs)^20^, was downregulated in Ror2-deficient basal cells, along with its paralog *Kif26a*, highlighting potential disruptions in cytoskeletal dynamics and cell fate regulatory mechanisms (**Figure 2H**).

### Wnt/Ror2 signaling maintains a basal/myoepithelial cell state

Gene expression analysis of p63^CreERT2/+^;ROSA^mTmG/+^ and p63^CreERT2/+^;ROSA^mTmG/+^;Ror2^fl/fl^ basal eGFP^+^ cells revealed widespread downregulation of basal cytokeratins following Ror2 deletion. After a 4-week chase period post-TAM treatment, WT mammary glands exhibited appropriate eGFP localization within the K14^+^ myoepithelial layer, with eGFP^+^/K14^+^ cells and K8^+^ cells remaining mutually exclusive within the epithelial bilayer (**Figure 3A**). As with previous studies, this observation suggests that basal cells in WT glands maintain their basal lineage identity over the 4-week chase period between puberty and adult development. In contrast, Ror2-deleted eGFP^+^ basal cells displayed a striking shift in localization, appearing within the K8^+^ luminal layer of the mammary gland (**Figure 3B**). These Ror2-deleted basal cells exhibit a loss of K14 expression and a concomitant gain of K8 expression, pointing to a lineage reprogramming within the epithelial bilayer upon Ror2 deletion. While Ror2-deleted basal cells lose basal cytokeratins and acquire luminal K8/18 expression within the inner luminal layer, they continued to express p63, suggesting a partial retention of basal features despite the loss of Ror2 (**Figure 3C, D**). Building on the lineage switch observed through immunostaining, we further validated this transition using flow cytometry to analyze surface markers CD24 and CD29, which distinguish luminal and basal/myoepithelial cell populations^7^. Under control conditions, eGFP^+^ basal cells exclusively comprised the basal compartment following a 4-week chase following TAM-mediated recombination (**Figure 3E-F**). However, following Ror2 deletion, approximately two-thirds of these basal cells shifted to a luminal cell fate within four weeks after TAM-mediated recombination (**Figure 3G**). Interestingly, while the overall distribution of hormone receptors within the luminal compartments remained unchanged between WT and Ror2-deleted conditions (**Figure 3H-J, L-N**), the majority of Ror2-KO basal cells adopted a luminal cell fate, with most expressing both Estrogen Receptor (ER) and Progesterone Receptor (PR) (**Figure H, I, K, L, M, N**). In addition to the observed shift toward ER^+^ and PR^+^ luminal cells, we examined GATA3 expression and found that the majority of eGFP^+^ Ror2-deleted cells within the luminal compartment were GATA3^+^ (**Figure S2**). This suggests that Ror2 loss not only drives a basal-to-luminal fate transition but also promotes differentiation into a GATA3^+^ luminal lineage, which is critical for maintaining luminal identity and function in the mammary gland.

**Figure 3.**
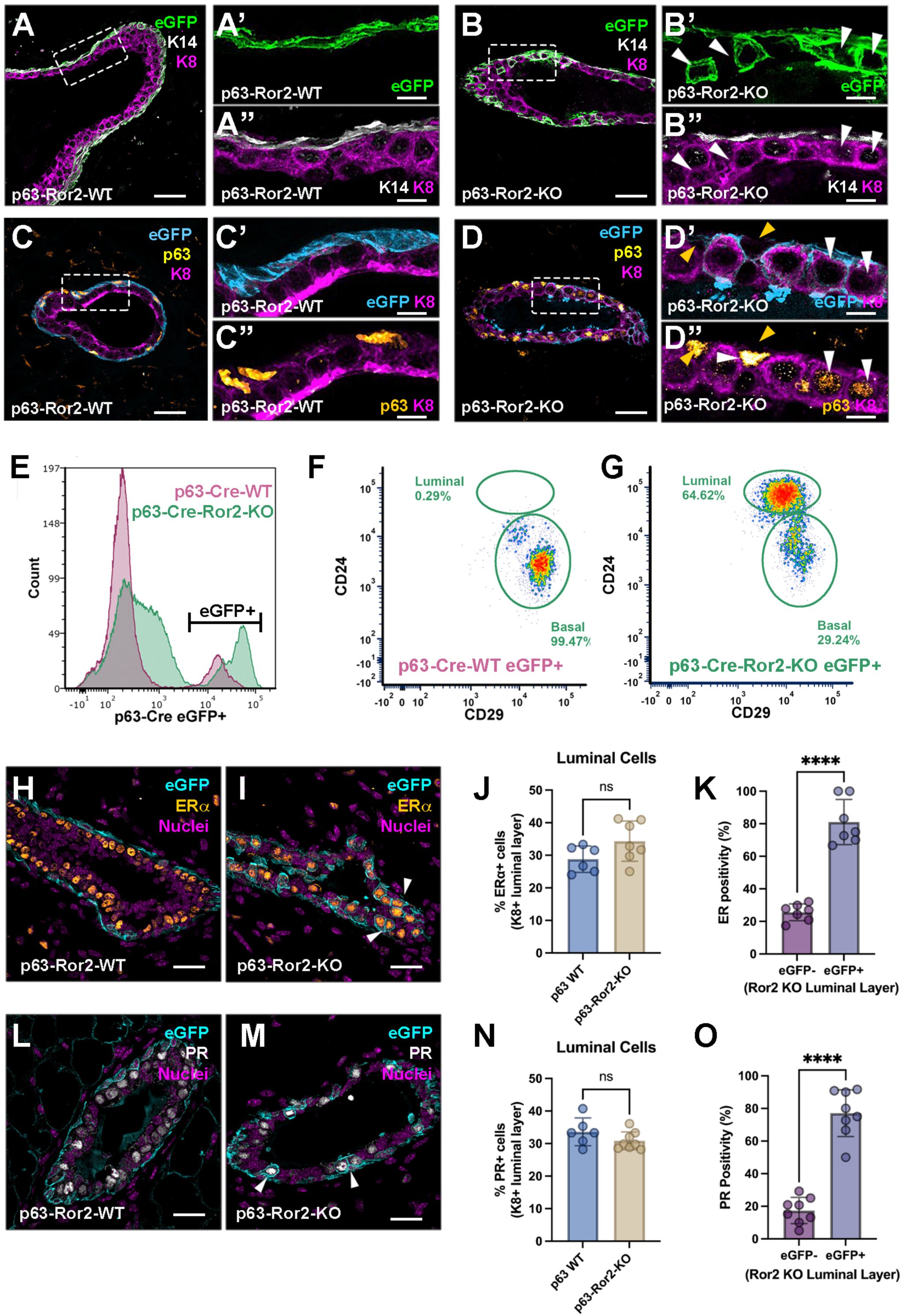
Ror2 loss in basal cells promotes cell plasticity and a fate transition into ER-positive luminal cells. (A, B) Representative immunofluorescence images showing K14 (gray), eGFP (green), and K8 (magenta) in mammary gland cross-sections from (A) control and (B) p63-Ror2-KO mice at 4 weeks post-tamoxifen injection. Scale bars: 25 μm for merged images and 10 μm for magnified insets (A’-B”). (B’, B”) White arrowheads highlight eGFP^+^ cells within the luminal compartment expressing K8 in p63-Ror2-KO mammary ducts. (C, D) Representative immunofluorescence images of p63 (yellow), eGFP (cyan), and K8 (magenta) in mammary gland cross-sections from (C) p63-Ror2-WT and (D) p63-Ror2-KO mice at 4 weeks post-tamoxifen injection. Scale bars: 25 μm for merged images and 10 μm for magnified insets (C’-D”). (D’-D”) White arrowheads indicate “hybrid” cells that are positive for both nuclear p63 and K8 in p63-Ror2-KO mammary ducts. Yellow arrows indicate p63^+^ basal epithelial cells. (E-F) Flow cytometry analysis (E) eGFP^+^ singlets from a pool of 3 WT (magenta) and 3 Ror2-KO glands (green) depicted by histogram. (F, G) CD24 and CD29 surface markers in eGFP^+^ recombined basal cells from (F) control and (G) p63-Ror2-KO mammary glands. A basal-to-luminal cell fate shift is observed in p63-Ror2-KO glands (10,000 single-cell events analyzed per group; representative of 3 independent experiments). (H, I) Representative immunofluorescence images showing eGFP (cyan) and ERα (orange) in mammary gland cross-sections from (H) control and (I) p63-Ror2-KO mice at 4 weeks post-tamoxifen injection. Scale bar: 20 μm. Nuclear ERα is detected in eGFP^+^ luminal cells within p63-Ror2-KO mammary ducts, while ERα is mutually exclusive from the eGFP^+^ basal cells in controls. (J) Quantification of the percentage of ERα^+^ luminal cells in control and p63-Ror2-KO groups (p = 0.0892; n = 6 and 7 random regions from three mammary glands, respectively). (K) Quantification of the percentage of ERα^+^ cells in eGFP^+^ and eGFP− luminal cells within the p63-Ror2-KO luminal layer (p < 0.0001; n = 7 random regions from three mammary glands). (L, M) Representative immunofluorescence images of eGFP (cyan) and PR (gray) in mammary gland cross-sections from (L) control and (M) p63-Ror2-KO mice at 4 weeks post-tamoxifen injection. Scale bar: 20 μm. Nuclear PR is detected in eGFP^+^ luminal cells within p63-Ror2-KO mammary ducts. (N) Quantification of the percentage of PR^+^ luminal cells in control and p63-Ror2-KO groups (p = 0.1677; n = 6 and 8 random regions from 3 mammary glands, respectively). (O) Quantification of the percentage of PR^+^ cells in eGFP^+^ and eGFP− luminal cells within the p63-Ror2-KO luminal layer (p < 0.0001; n = 8 random regions from 3 mammary glands).

### Ror2 Loss in the basal compartment drives lineage-specific chromatin remodeling

Given the observed lineage changes upon Ror2 loss, we sought to investigate whether these shifts were associated with alterations in chromatin accessibility at lineage-specific regulatory regions. To address this, we performed single-cell ATAC sequencing (scATAC-seq) on FACS-sorted eGFP^+^ cells isolated from control (p63^CreERT2/+^;ROSA^mTmG/+^) and experimental (p63^CreERT2/+^;ROSA^mTmG/+^;Ror2^fl/fl^) mice two weeks after TAM-mediated recombination. This approach allowed us to interrogate chromatin accessibility dynamics within basal and luminal epithelial compartments and identify potential regulatory changes underlying the basal-to-luminal cell fate transition following loss of Wnt/Ror2 signaling *in vivo*. UMAP projections of scATAC-seq data revealed distinct clustering patterns between basal and luminal epithelial subsets, providing insight into chromatin accessibility changes associated with Ror2 loss (**Figure 4A**). In control p63^CreERT2/+^;ROSA^mTmG/+^ eGFP^+^ basal cells, a single, well-defined cluster was observed, indicative of a uniform basal population. In contrast, eGFP^+^ cells from experimental p63^CreERT2/+^;ROSA^mTmG/+;Ror2fl/fl^ mice formed three distinct clusters, none of which fully overlapped with the WT basal population. Analysis of UMAP projections for lineage markers such as *TP63* (**Figure 4B**) and *ESR1* (Estrogen Receptor 1) (**Figure 4C**) revealed distinct basal and luminal subpopulations within the experimental condition, highlighting a reorganization of lineage-specific chromatin accessibility upon Ror2 loss. These changes corresponded to a shift in basal cells toward luminal characteristics, with the emergence of heterogeneous subpopulations. A volcano plot of differentially accessible genes highlighted significant upregulation of *ESR1*, *ESR2*, *MAFA*, and *PPARG* in Ror2-deficient clusters further illustrates these changes in the regulatory landscape (**Figure 4D**). These findings suggest that Ror2 loss promotes the activation of luminal-associated chromatin accessibility and regulatory programs. Conversely, genes linked to basal identity, including *TP63, SOX12, GLIS3, EGR3, CENPB,* and *ZFP57*, were significantly downregulated (**Figure 4D**). The increased expression of *MAFA* and *PPARG*, transcription factors associated with differentiation and metabolic programming, further supports a basal-to-luminal transition at the chromatin level. These results collectively suggest that Ror2 loss disrupts basal cell identity and facilitates the acquisition of luminal-like features within the epithelial compartment.

**Figure 4.**
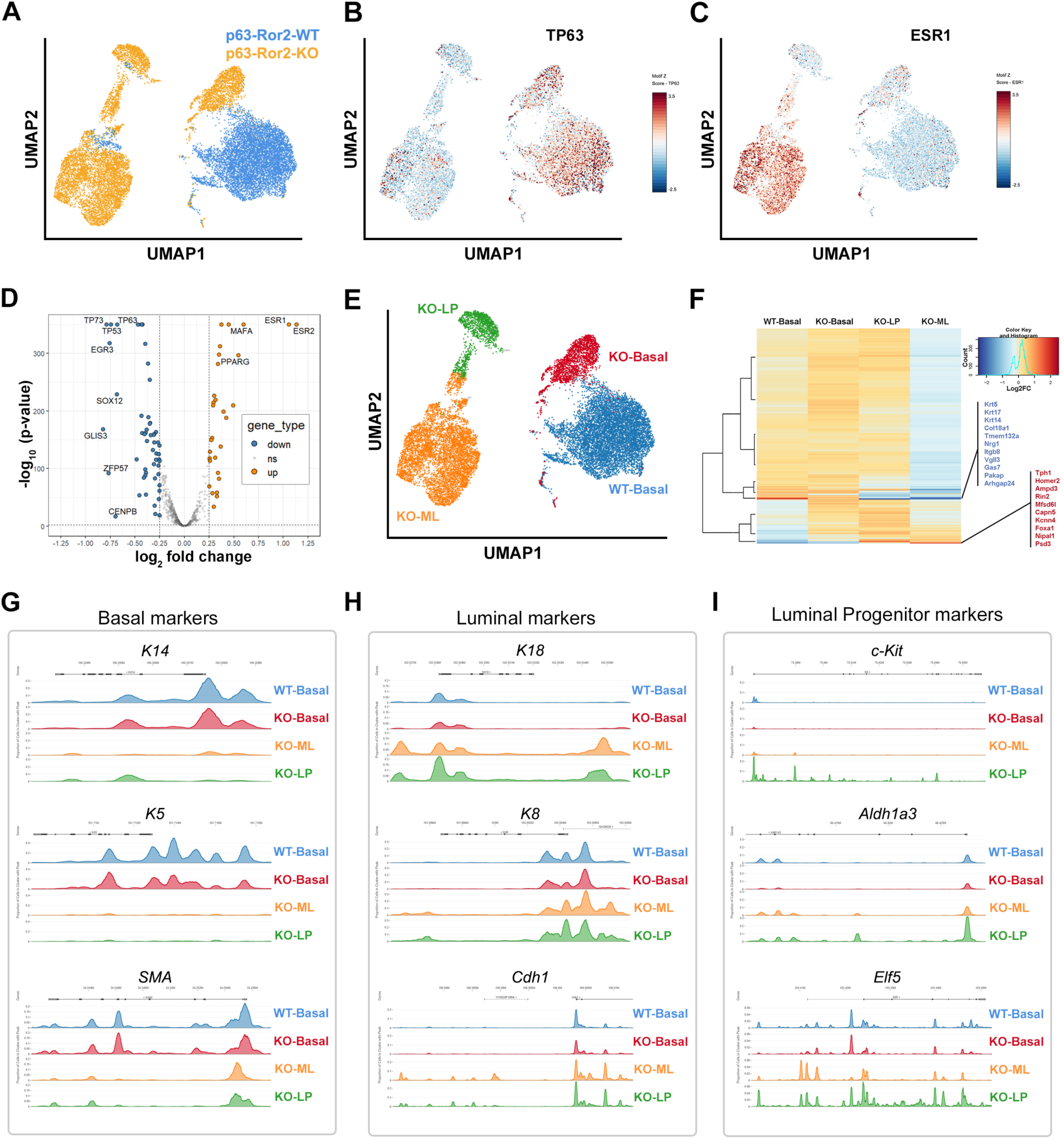
Single-cell ATAC-sequencing reveals transcriptional and chromatin changes induced by Ror2 loss in basal mammary cells. (A) Uniform manifold approximation and projection (UMAP) visualization of single-cell ATAC-seq data, showing 8,823 control (blue) and 9,500 p63-Ror2-KO (yellow) eGFP^+^ mammary epithelial cells. (B, C) Heatmaps displaying Z-scores of transcription factor motif enrichment for (B) TP63 and (C) ESR1 across single cells in the UMAP including both control and p63-Ror2-KO eGFP^+^ cells. Enrichment scores are represented by a gradient from blue (low score) to maroon (high score). (D) Volcano plot illustrating significantly upregulated (orange; adjusted p < 0.05 and log₂FC > 0.25) and downregulated (blue; adjusted p < 0.05 and log₂FC < –0.25) motifs in p63-Ror2-KO cells compared to p63-WT control cells. Motifs not significantly altered are shown in gray. (E) UMAP visualization of scATAC-seq data colored by cell type. Control cells primarily cluster as basal cells (blue), while p63-Ror2-KO cells distribute into basal (red), luminal progenitor (green), and ER^+^ luminal (orange) cell types. (F) Heatmap showing significantly differentially expressed genes (adjusted p < 0.05) identified through promoter enrichment analysis among four cell types. Expression levels are represented as log_2_FC using a gradient from blue (downregulation) to maroon (upregulation). Genes within the two most significantly altered clusters are listed. (G) Peak tracks depicting accessible genomic regions for basal marker genes, including *Keratin 14*, *Keratin 5*, and s*mooth muscle actin*. These genomic elements show greater accessibility in basal cell types in both control and p63-Ror2-KO groups. (H) Peak tracks showing accessible genomic elements for luminal marker genes, including *Keratin 18*, *Keratin 8*, and *Cdh1* (*E-cadherin*). These regions exhibit increased accessibility in luminal and luminal progenitor cell types in the p63-Ror2-KO eGFP^+^ cells. (I) Peak tracks highlighting accessible genomic regions for luminal progenitor marker genes, including *c-Kit, Aldh1a3*, and *Elf5*. These regions are more accessible in luminal progenitor cell types in the p63-Ror2-KO eGFP^+^ cells.

To further delineate the impact of Ror2 loss on basal-to-luminal transitions, we utilized publicly available datasets to refine UMAP1 vs. UMAP2 clustering based on key genes known to be involved in mammary epithelial lineages^24,25^. In control p63^CreERT2/+^;ROSA^mTmG/+^ (WT) eGFP^+^ cells, a single distinct basal cluster was identified. In contrast, Ror2-deficient p63^CreERT2/+^;ROSA^mTmG/+^;^Ror2fl/fl^ eGFP^+^ cells formed three distinct clusters, corresponding to Ror2-KO basal, luminal progenitor (LP), and mature luminal (ML) populations (**Figure 4E**). Hierarchical clustering of gene expression profiles visualized via heatmap highlighted similarities in basal-specific gene expression between WT and KO basal populations (**Figure 4F**). However, Ror2-KO basal cells exhibited notable upregulation of luminal-specific genes, including *Foxa1*, *Tph3*, *Homer2*, *Ampd3*, *Rin2*, *Capn5*, and *Psd3*, suggesting a partial acquisition of luminal characteristics (**Figure 4F**). This luminal shift was further substantiated by the loss of basal-specific genes such as *Krt5*, *Krt14*, *Krt17*, *Nrg1*, and *Vgll3* in Ror2-KO basal cells. Differential chromatin accessibility analyses revealed loss of regulatory peaks associated with *K14*, *K5*, and *SMA* in luminal progenitor and mature luminal populations after Ror2 loss (**Figure 4G**). Conversely, luminal markers *K8*, *K18*, and *Cdh1* were gained (**Figure 4H**), signifying a chromatin landscape favoring luminal differentiation. Moreover, luminal progenitor-specific markers, including *c-kit*, *Aldh1a3*, and *Elf5*, were upregulated, providing further resolution between the LP and ML clusters (**Figure 4I**). Together, these data highlight a dramatic reorganization of chromatin accessibility and gene expression upon Ror2 loss in basal cells, underscoring the role of Ror2 in maintaining basal lineage integrity and preventing luminal lineage acquisition.

### Ror2 loss disrupts RhoA-ROCK1 and YAP1 signaling to drive basal-to-luminal transcriptional reprogramming

We began by performing Reactome pathway analysis on genes significantly downregulated in Ror2-KO basal populations compared to WT. This analysis identified top pathways, including signaling by Rho GTPases, CDC42 GTPase signaling, axon guidance, and ephrin signaling, all of which are critical for cell-cell communication and actin cytoskeletal dynamics (**Figure 5A**). These findings align with the established role of noncanonical Wnt signaling, mediated by Ror2, in regulating cytoskeletal organization. Focusing on Rho signaling, we confirmed the downregulation of *Arhgap32* and *Arhgap24*, key regulators of Rho GTPase activity, in p63Cre-Ror2-KO eGFP^+^ basal cells using qRT-PCR (**Figure 5B**). To validate this pathway *in vitro*, we utilized lentiviral shRNA-mediated depletion of Ror2 in basal epithelial cells and observed a significant reduction in RhoA-GTP levels, as assessed by G-LISA, compared to shLUC controls (**Figure 5C, D**). This loss of active RhoA was accompanied by reduced total protein levels of RhoA and ROCK1, confirmed by Western blotting (**Figure 5D**). To further investigate the role of Ror2 in activating RhoA, we treated cells with Wnt5a, a known ligand for Ror2. In shLUC control cells, Wnt5a treatment significantly increased RhoA-GTP levels; however, this activation was completely abrogated in Ror2-depleted cells (**Figure 5D**). Importantly, no effects on CDC42 or Rac1/2/3 activity were observed, indicating a selective role for Ror2 in RhoA-mediated signaling downstream of Wnt5a. These findings suggest that Ror2 specifically modulates RhoA signaling to maintain cytoskeletal dynamics and basal cell integrity.

**Figure 5.**
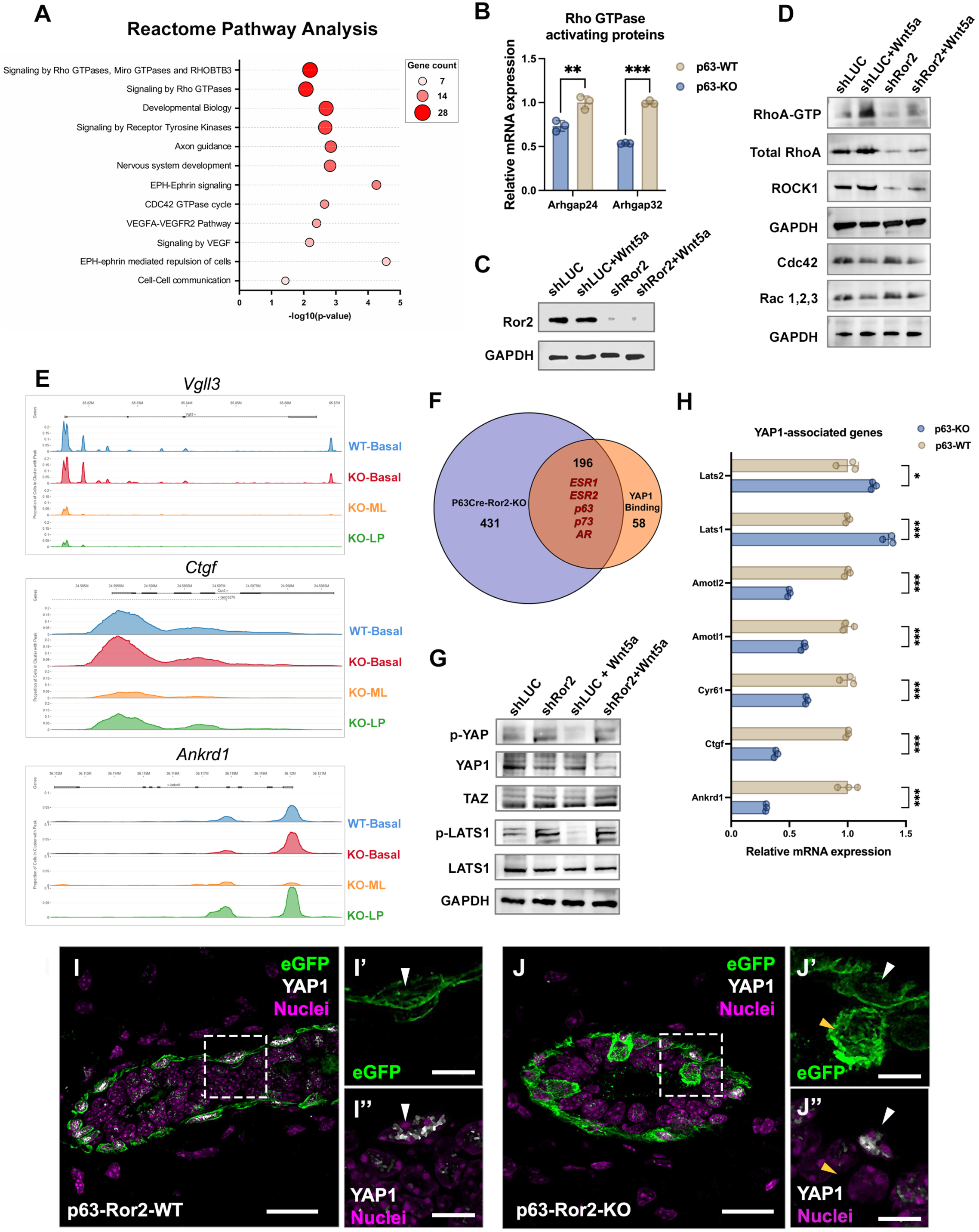
Ror2 loss impairs RhoA activity and YAP1 signaling. (A) Bubble plot showing Reactome Pathway Analysis enrichment based on genes with significantly altered promoter accessibility in scATAC-seq data from control and p63-Ror2-KO cells. Pathways are ranked by the number of genes with altered promoter accessibility, represented by bubble size and red color gradients. (B) Quantitative RT-qPCR analysis of Rho GTPase-activating proteins *Arhgap24* and *Arhgap32* in FACS-isolated basal epithelial cells from control and p63-Ror2-KO mice. Gene expression levels were normalized to GAPDH, and fold changes were plotted relative to the control group (*p* = 0.008 for *Arhgap24*; *p* < 0.001 for *Arhgap32*; *n* = 3 biological replicates per group). (C) Western blot for Ror2 showing efficient Ror2 knockdown with LeGO-shRor2 cells relative to LeGO-shLUC control cells. (D) Western blot analysis of GTP-bound RhoA, total RhoA, ROCK1, Cdc42, Rac1/2/3, and GAPDH in shLUC and shRor2 mammary basal epithelial cells with or without Wnt5a treatment. (E) Peak tracks illustrating accessible genomic elements for YAP1 downstream target genes Vgll3, Ctgf, and Ankrd1. These regions are less accessible in luminal cell types in p63-Ror2-KO groups. (F) Venn diagram showing the overlap between significantly altered transcription factors identified in scATAC-seq motif enrichment analysis and YAP1-binding transcription factors from previous ChIP sequencing data^26^. (G) Western blot analysis of phospho-YAP1, total YAP1, TAZ, phospho-LATS1, LATS1, and GAPDH in shLUC and shRor2 mammary basal epithelial cells with or without Wnt5a treatment. (H) Quantitative RT-qPCR analysis of Hippo/YAP1 signaling genes in FACS-isolated basal epithelial cells from control and p63-Ror2-KO mice. Gene expression levels were normalized to *GAPDH*, and fold changes were plotted relative to the control group (*p* = 0.015 for LATS2; *p* < 0.001 for other genes; *n* = 3 biological replicates per group). (I, J) Representative immunofluorescence images showing eGFP (green) and YAP1 (gray) in mammary gland cross-sections from (I) control and (J) p63-Ror2-KO mice at 4 weeks post-tamoxifen injection. Scale bars: 20 μm for merged images and 10 μm for magnified insets (I’, I”, J’, J”). White arrowheads indicate nuclear YAP1 in eGFP^+^ basal cells, while yellow arrowheads indicate the absence of nuclear YAP1 in eGFP^+^ luminal cells within p63-Ror2-KO mammary ducts.

Given the known role of RhoA-ROCK1 signaling in regulating mechanosensitive pathways, we next investigated the impact of Ror2 loss on YAP1-associated signaling, which controls focal adhesion integrity downstream of RhoA. scATAC-seq analysis revealed reduced chromatin accessibility at loci for YAP1 target genes, including Vgll3, Ctgf, and Ankrd1, in p63Cre-Ror2-KO eGFP^+^ basal cells, suggesting impaired YAP1 transcriptional activity (**Figure 5E**). By integrating scATAC-seq data with transcription factor motif analyses^26^, we identified significant enrichment for motifs corresponding to *ESR1, ESR2, p63, p73*, and *AR* at Ror2-regulated sites (**Figure 5F**). Notably, p73, a well-established YAP cofactor, and ESR1/ESR2, transcription factors not previously linked to YAP activity in this context, emerged as potential mediators of chromatin and transcriptional changes upon Ror2 loss (**Figure 5F**). To further validate these findings, we demonstrated that Wnt5a treatment represses phosphorylation of YAP1 (pYAP1) in shLUC control cells, consistent with YAP1 activation (**Figure 5G**). However, this repression was abrogated upon Ror2 depletion, indicating a critical role for Ror2 in mediating Wnt5a-dependent YAP1 activation (**Figure 5G**). Moreover, while Wnt5a treatment reduced phosphorylation of LATS1 in shLUC cells, this effect was impaired in Ror2-depleted cells. Consistent with these results, p-LATS1 levels were significantly elevated in p63Cre-Ror2-KO basal cells, correlating with a reduction in active YAP1 signaling (**Figure 5G**). qRT-PCR confirmed these findings, showing increased expression of Lats1/2 and downregulation of YAP1 target genes^27–29^, including *Amotl1*, *Amotl2*, *Cyr61*, *Ctgf*, and *Ankrd1*, in Ror2-deficient basal cells (**Figure 5H**). The downregulation of YAP1 was confirmed *in vivo*, where eGFP^+^ Ror2-KO basal cells that shifted into the luminal compartment upon Ror2 loss and luminal differentiation exhibited a loss of YAP1 expression (**Figure 5I, J**). Together, these data reveal a critical link between Ror2-mediated RhoA signaling and YAP1 activation. The involvement of motifs for p73 and nuclear receptors like ESR1 and ESR2 further suggests a cooperative network of transcription factors contributing to the basal-to-luminal shift observed upon Ror2 loss.

### RhoA-ROCK1 activity is essential to maintain basal cell identity

To further investigate the role of RhoA-ROCK1 signaling downstream of Ror2 in maintaining basal cell identity, we employed an organoid-based approach to assess epithelial lineage maintenance under conditions of ROCK1 inhibition. This approach followed up on our earlier findings that demonstrated impaired RhoA activation and reduced ROCK1 levels in Ror2-deficient basal cells (**Figure 5D**). Using organoids isolated from p63^CreERT2/+^;ROSA^mTmG/+^ mammary glands after Cre-mediated recombination and lineage labeling of the basal compartment, we tested whether ROCK1 inhibition was sufficient to drive a basal-to-luminal shift. In vehicle-treated organoids, eGFP^+^ basal cells remained confined to the basal layer at the periphery of the organoid and were mutually exclusive from luminal ER^+^ and K8 expression (**Figure 6A, D-E**). Similar to basal-specific Ror2 deletion (**Figure 6B**), treatment with the selective ROCK inhibitor Y27632 led to a significant loss of basal cell identity, as evidenced by the acquisition of luminal markers ER and K8 in eGFP-labeled p63^+^ basal cells (**Figure 6C, D-E**). This phenotypic switch was observed within a 4-day chase period *in vitro* under Y27632 treatment, indicating that disruption of ROCK1 activity is sufficient to reprogram basal cells toward a luminal cell fate.

**Figure 6.**
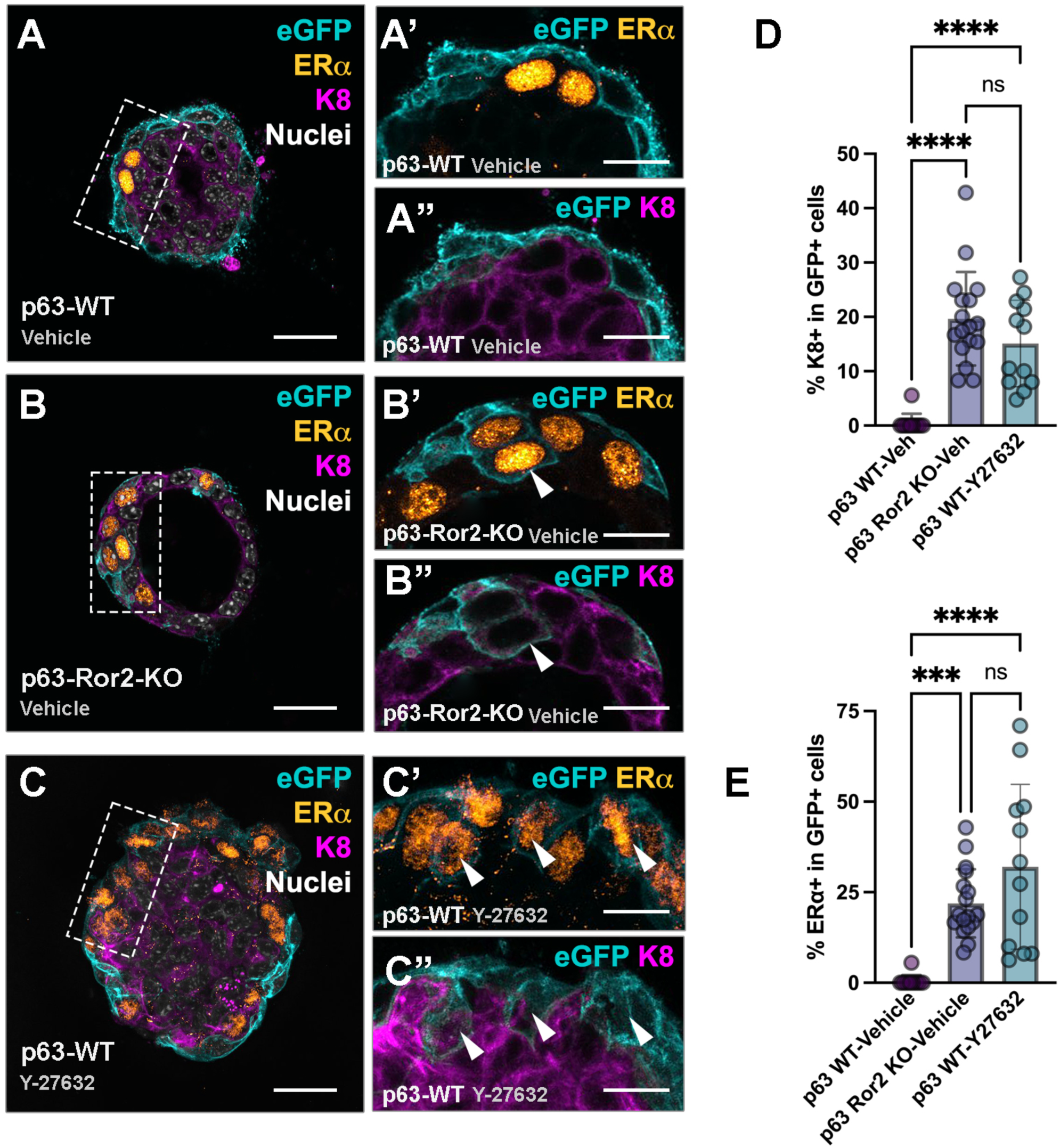
ROCK inhibition induces ERα and K8 expression in basal cells. (A-C) Representative immunofluorescence images showing eGFP (cyan), ERα (orange), K8 (magenta), and nuclei (gray) in cross-sections of mammary gland organoids cultured in Matrigel for 4 days under three conditions: (A) control WT, (B) p63-Ror2-KO, and (C) WT treated with the ROCK inhibitor Y-27632. (A’-C“) Insets within A-C showing eGFP (cyan) with ERα (orange) or eGFP (cyan) with K8 (magenta). Vehicle controls exhibit mutual exclusivity between eGFP and ERα, as well as eGFP and K8, unlike Y-27632 or Ror2-KO organoids. Scale bar: 20 μm. White arrowheads indicate eGFP^+^ cells expressing ERα. (D) Quantification of the percentage of ERα+ cells among eGFP^+^ cells across the 3 groups. Statistical comparisons: *p* = 0.0003 for WT-Veh vs. KO-Veh; *p* < 0.0001 for WT-Veh vs. WT-ROCKi; *p* = 0.06 for KO-Veh vs. WT-ROCKi (*n* = 11, 17, and 12 random organoids from 3 independent wells, respectively). (E) Quantification of the percentage of K8^+^ cells among eGFP^+^ cells in the three groups. Statistical comparisons: *p* < 0.0001 for WT-Veh vs. KO-Veh; *p* < 0.0001 for WT-Veh vs. WT-ROCKi Y-27632; *p* = 0.10 for KO-Veh vs. WT-ROCKi Y-27632 (*n* = 11, 17, and 12 random organoids from three independent wells, respectively).

These findings provide direct evidence that RhoA-ROCK1 signaling downstream of Ror2 is critical for preserving basal cell state and preventing luminal lineage acquisition. The ability of ROCK1 inhibition to phenocopy the effects of Ror2 loss further underscores the mechanistic link between Ror2 signaling and the maintenance of basal epithelial identity (**Figure 6B-C, D-E**).

## Discussion

In this study, we demonstrated that Ror2, a receptor central to noncanonical Wnt signaling, plays a pivotal role in maintaining basal epithelial cell identity and regulating branching morphogenesis during mammary gland development. Using a p63Cre^ERT2/+^ lineage-specific model, we achieved efficient and targeted deletion of Ror2 within the basal epithelial compartment. Our findings reveal that the basal-specific deletion of Ror2 disrupts both morphogenic processes and cellular fate programs, underscoring its critical role in coordinating the structural and identity-related aspects of mammary gland development. While primary ductal extension remained unchanged, secondary and tertiary branching were markedly enhanced, suggesting that Wnt/Ror2 signaling in the basal epithelial layer constrains the plasticity required for higher-order morphogenetic processes, akin to the role of TGF-β^30–32^. Indeed, multiple studies have shown that mammary branching constraint is largely dictated by TGF-β, with downstream effectors such as Wnt5a^33^ and Slit/Robo^34^ implicated in this spatial patterning. Prior studies, including our own, demonstrated that Ror2 loss-of-function in the mammary epithelium enhances branching *in vivo*^21^; however, the specific cellular compartment responsible for this effect remained undefined until now. Our findings highlight the predominant role of Wnt/Ror2 signaling within the basal/myoepithelial compartment in driving this phenotype.

To investigate transcriptional changes within basal cells upon Ror2 loss, we uncovered an unexpected and striking finding: Ror2-KO basal cells exhibited dramatic alterations in lineage-specific markers, suggesting that more than morphogenetic regulation was at play. We identified a basal-to-luminal transition marked by the downregulation of basal markers, including Keratins 5, 14, and 17, and the upregulation of luminal markers such as Keratins 8 and 18. Lineage tracing and gene expression profiling revealed this shift in lineage identity, while single-cell chromatin accessibility analyses uncovered a profound reorganization of the transcriptional and epigenetic landscape, driving the transition from the basal lineage to luminal-restricted state. These findings implicate impaired Ror2 signaling as a driver of a basal-to-luminal lineage transition specifically under normal physiological contexts and during postnatal mammary gland ductal morphogenesis. Importantly, our results distinguish the role of Ror2 from previously reported settings where basal cells adopt a luminal cell state in response to cellular stresses. For instance, basal-to-luminal transitions have been observed under genotoxic stress induced by DNA-damaging agents such as cisplatin and MMC (DNA crosslinking agents)^35^, in *ex vivo* dissociation and transplantation settings (e.g., into the cleared mammary fat pad)^7,8,10,36^, or following luminal cell ablation^37^. In these contexts, basal cells are typically provoked into a luminal fate through reactivation of a multipotency program. Our findings identify a novel role for Ror2 signaling in maintaining basal cell identity under homeostatic conditions, independent of external stressors or perturbations, suggesting that noncanonical Wnt signaling preserves basal cell identity *in situ*.

Mechanistically, we demonstrate that Ror2 regulates basal cell integrity via the RhoA-ROCK1-YAP1 axis. Loss of Ror2 reduces RhoA and ROCK1 activity, resulting in a disorganized actin cytoskeleton and attenuated YAP1 signaling. Previous studies have also suggested a role for miR205 in the regulation of the YAP pathway in mammary basal cells^38^. This disruption plausibly links alterations in cellular mechanics to distinct cell fate decisions between basal and luminal compartments, a phenomenon similarly observed in other epithelial systems where YAP signaling integrates mechanical cues to regulate lineage fidelity and differentiation^39^. These disruptions were accompanied by changes in chromatin accessibility at basal– and luminal-specific loci, implicating transcriptional and epigenetic regulators—such as p63, p73, ESR1, and ESR2—in mediating the basal-to-luminal cell fate switch. Inhibition of ROCK1 in organoids phenocopied the effects of Ror2 deletion, providing compelling evidence that the RhoA-ROCK1 signaling axis is essential for maintaining basal cell identity. Studies in embryonic stem cells have shown that ROCK inhibition supports cell survival *in vitro*^40,41^, while in mesenchymal stem cells, it influences lineage fate through regulation of a CTCF-VEGF complex within the extracellular matrix^42^. Similarly, prior studies with mammary basal cells in feeder-layer adherent colonies demonstrated that actin cytoskeleton remodeling and TGF-β downregulation enhance colony-forming efficiency^41^. Extending previous reports that ROCK1 inhibition results in abnormal branching morphogenesis^43^—where its inhibition via Y-27632 led to disorganized branching structures and inappropriate myoepithelial cell coverage—we now demonstrate that ROCK1 disruption drives p63^CreERT2/β^;ROSA^mTmG/+^ eGFP^+^ basal cells to acquire ER^+^ and K8^+^ luminal cell features in organoid cultures. Together, these findings underscore how impaired Ror2 signaling destabilizes basal cell integrity, driving basal-to-luminal transitions through disruption of cytoskeletal dynamics, altered YAP1 signaling, and reprogramming of chromatin accessibility. Known roles for Notch signaling in the establishment and maintenance of luminal cell fate suggests that Wnt/Ror2 and Notch pathways likely coordinate to regulate the balance between basal and luminal cell states^44–46^. Notch1 also plays a critical role in designating alveolar progenitors necessary for pregnancy^44^, raising further questions about how Wnt/Ror2 signaling intersects with Notch to maintain cell identity and function during other stages, such as pregnancy and lactation. This interplay highlights the broader significance of Ror2 and ROCK1 as key regulators that integrate cytoskeletal dynamics with lineage-specific transcriptional programs to preserve basal cell identity and developmental fidelity in the mammary gland. Future studies exploring the crosstalk between Wnt/Ror2 and Notch signaling pathways could provide deeper insights into their cooperative roles in mammary gland biology.

Tumor cell heterogeneity in cancer biology is frequently linked to lineage infidelity, cell fate plasticity, and the acquisition of multipotency, emphasizing the critical need to understand cell state fidelity in normal development to elucidate its role in cancer progression. In triple-negative breast cancers (TNBC), basal-like TNBC has been linked to the luminal progenitor as the cell of origin^47,48^. Transcription factors such as Sox9 have been identified as critical determinants driving the transition from a luminal progenitor to a basal-like cell state, initiating basal-like breast cancers *in situ*^49^. Similarly, oncogenic activation of PIK3CA has been shown to trigger a luminal-to-basal cell fate transition and promote basal-like TNBC in mouse models^50^. These observations underscore how deviations in signaling pathways, environmental cues, or hormonal alterations can disrupt developmental programs, skewing lineage fidelity and driving the acquisition of altered cellular fates *in situ*.

Collectively, our results establish Ror2 as a critical regulator of basal epithelial identity and function, with broad implications for understanding mammary gland morphogenesis and lineage plasticity. The observed basal-to-luminal transition upon Ror2 loss underscores the plasticity of the basal compartment and highlights how disruptions in signaling networks can alter epithelial hierarchies. These findings may offer valuable insights into the mechanisms driving lineage plasticity in breast pathologies, including tumor cell transitions and the development of cellular heterogeneity in breast cancer.

## Figure Legends

**Figure S1.** Loss of Ror2 disrupts the expression of genes associated with cytokeratin, cell junctions, adhesion, and extracellular matrix (ECM) remodeling. Quantitative RT-qPCR analysis of relative mRNA expression for genes involved in cytokeratin organization (*K14, K5, K6a*), cell junction and adhesion (*Cldn1, Bcam1, Robo1, Robo2, Flrt2, Dsg1a, Dsg1b, Dsg1c, Dsg3, Dsc3, Kif26b*), and ECM regulation (*Fn1*) in mammary epithelial cells from p63-Ror2-WT (gray) and p63-Ror2-KO (orange) mice. Gene expression levels were normalized to *GAPDH*, and data are presented as fold change relative to the control group. Statistical significance is denoted by *p<0.05, **p<0.01, and ***p<0.001. n=3 biological replicates per group.

**Figure S2.** Loss of Ror2 in basal cells drives a shift towards luminal cell identity. (A-B) Representative confocal images of mammary ducts from (A) p63-Ror2-WT (B) p63-Ror2-KO mice at 6 weeks old immunostained for estrogen receptor alpha (ERα, magenta), eGFP (cyan), and nuclei (gray). Scale bar, 50 μm. (C-D) Quantification of the percentage of ERα^+^ cells among all luminal cells in (C) WT and Ror2-KO mice (p = 0.92; n = 5 and 10 random regions from three mammary glands, respectively) and (D) among eGFP^+^ and eGFP^−^ luminal cells in Ror2-KO mice (p < 0.0001; n = 10 and 10 random regions from three mammary glands). Statistical significance: ns = not significant, ****p<0.0001. (E-F) Representative confocal images of mammary ducts from (E) p63-Ror2-WT and (F) p63-Ror2-KO mice immunostained for progesterone receptor (PR, magenta), eGFP (cyan), and nuclei (gray). Scale bars, 50 μm. 6 weeks of age (G-H) Quantification of the percentage of PR+ cells among all luminal cells in WT and Ror2-KO mice (p = 0.85; n = 7 and 6 random regions from three mammary glands, respectively). (G) and among eGFP^+^ and eGFP− luminal cells in Ror2-KO mice (p = 0.0004; n = 6 and 6 random regions from three mammary glands). (H). Statistical significance: ns = not significant, ***p<0.001. 6 weeks of age (I-J) Representative confocal images of mammary ducts from p63-Ror2-WT (I) and p63-Ror2-KO (J) mice immunostained for GATA3 (magenta), eGFP (cyan), and nuclei (gray). Scale bars, 20 μm. 6 weeks of age.

## Methods

### Animal studies

This study adhered to the *Guide for the Care and Use of Laboratory Animals* of the National Institutes of Health and was approved by the Baylor College of Medicine Institutional Animal Care and Use Committee. To investigate the role of Ror2 in the mammary epithelium, we generated a myoepithelial-specific Ror2 conditional knockout model using a previously generated knock-in lineage-specific p63^CreERT2^ mice crossed with Ror2^f/f^ conditional mutant mice, obtained from Jackson Laboratories^20^. *p63^CreERT^*^2^ knock-in mice were obtained from Dr. Jianming Xu and generated as described previously^51^. Male p63^CreERT2/+;^ Ror2^f/wt^ mice were bred with homozygous ROSA^mTmG^ double-fluorescent reporter females to produce ROSA^mTmG/+^; p63^CreERT2/+^; Ror2^wt/wt^ males. These males were crossed with wild-type females to establish the control cohort (ROSA^mTmG/+^; p63^CreERT2/+^; Ror2^wt/wt^). For the experimental cohort, ROSA^mTmG/+^; p63^CreERT2/+^; Ror2^f/wt^ males were bred with Ror2^f/f^ females to generate ROSA^mTmG/+^; p63^CreERT2/+^; Ror2^f/f^ males, which were subsequently crossed with Ror2^f/f^ females to yield myoepithelial-specific Ror2 homozygous floxed female mice (ROSA^mTmG/+^; p63^CreERT2/+^; Ror2^f/f^). All mouse lines were maintained on a C57Bl/6 genetic background. Genotyping of all mice was performed using PCR-based methods as described^20^. Cre^ERT2^ recombinase activity was induced at 4 weeks of age for pubertal studies or at 6 weeks of age for adult virgin studies. Tamoxifen (TAM; Sigma-Aldrich, #T5648) was prepared at a concentration of 20 mg/ml in a solution of 90% corn oil and 10% ethanol. To achieve efficient recombination, mice received three consecutive intraperitoneal injections of TAM at a dose of 80 mg/kg body weight. This TAM regimen has been shown to have no observable impact on normal mammary epithelial development^52^. Tissues were harvested at two distinct time points: 48 hours post-TAM administration for initial labeling studies and 2-or 4-weeks post-TAM for lineage tracing and functional analyses. These time points were chosen to capture both the immediate and long-term effects of Ror2 deletion in the myoepithelial compartment.

### Preparation of Primary Organoids

Primary mammary organoids were isolated from 8-to 12-week-old female mice. Mammary glands were dissected and minced into ∼1 mm³ fragments. The tissue was enzymatically dissociated in Hanks’ Balanced Salt Solution (HBSS) containing 1 mg/ml collagenase A (Roche, #11088793001) and 1 μg/ml DNase I (StemCell Technologies, #07900) in a shaking incubator at 37°C, rotating at 125 rpm for 2 hours. During digestion, the mixture was gently pipetted every 20 minutes to ensure uniform dissociation. To enrich for mammary epithelial organoids, the digested tissue was subjected to differential centrifugation at 1,500 rpm for 10 seconds, repeated three times. The resulting organoids were either embedded in growth factor-reduced matrices for branching morphogenesis assays or further processed. For downstream analyses, such as flow cytometry, colony-forming assays, and mammosphere assays, the organoids were digested into single cells using 0.25% Trypsin-EDTA (ThermoFisher Scientific, #25200056).

### Branching Morphogenesis Assays

Twenty-four-well culture plates (Corning, #3524; Corning, NY, USA) or eight-chamber slides (Corning, #354118) were pre-coated with 5 μl of growth factor-reduced Matrigel (Bio-Techne, #3536-005-02; Minneapolis, MN, USA) and incubated at 37°C for 15 minutes to allow for polymerization. Freshly isolated primary mammary organoids were resuspended in Matrigel at a concentration of approximately 1000 organoids/ml. A 40 μl aliquot of the organoid suspension was plated into each well of the 24-well plates or chamber of the eight-chamber slides. The plated cultures were incubated at 37°C for 1 hour to permit gel polymerization. Following polymerization, Matrigel layers were overlaid with 750 μl of growth media in 24-well plates or 500 μl in eight-chamber slides. The growth media consisted of DMEM/F12 (ThermoFisher Scientific, #11320033; Waltham, MA, USA) supplemented with 1% insulin-transferrin-selenium (ITS Liquid Media Supplement, Sigma-Aldrich, #I3146; St. Louis, MO, USA), 1% penicillin/streptomycin (ThermoFisher Scientific, #15140-122), and 50 ng/ml FGF2 (Sigma-Aldrich, #F0291). Media containing growth factors were replaced every other day. To assess morphological changes during organoid development, cultures were photographed daily from days 3 to 6 using brightfield microscopy.

### Tissue Processing and Histology

To assess cell proliferation, 5-bromo-2′-deoxyuridine (BrdU; 60 μg/g body weight) was administered to mice via intraperitoneal injection 2 hours prior to tissue collection. Mammary fat pads were dissected and subjected to fluorescent whole-mount imaging. For imaging, mammary glands were gently sandwiched between two glass slides and visualized using a stereomicroscope (Leica) to capture ductal architecture and fluorescent labeling.

Following imaging, mammary glands were fixed in 4% paraformaldehyde (PFA) at 4°C overnight to preserve tissue structure. After fixation, tissues were processed into paraffin blocks for histological analysis. Organoid cultures were similarly fixed by washing in phosphate-buffered saline (PBS) and incubating in 4% PFA for 10 minutes at room temperature. To ensure the integrity and orientation of 3D organoids embedded in Matrigel, samples were further embedded in HistoGel (Epredia, #22-11-678) before paraffin processing. Mammary glands or organoids embedded in paraffin were sectioned into 5-μm-thick slices using a microtome. Sections were prepared for subsequent immunostaining and histological analyses. For wholemount confocal imaging of mammary glands, CUBIC (Clear, Unobstructed Brain/Body Imaging Cocktails and Computational Analysis) clearing was carried out as previously described^22,23^.

### Immunofluorescence Staining of Tissue and 3D Organoid Sections

Tissue and organoid sections were prepared for immunofluorescence staining through deparaffinization and rehydration. Sections were deparaffinized using xylene washes followed by a graded ethanol series (100%, 95%, 70%) and rehydrated in distilled water. Heat-induced epitope retrieval was performed using sodium citrate buffer (10 mM sodium citrate, pH 6.0) or Tris-EDTA buffer (10 mM Tris, 1 mM EDTA, pH 9.0) at 95°C for 20 minutes in a microwave. Sections were then cooled to room temperature for 30 minutes and rinsed thoroughly with 1× PBS twice for 10 minutes. Blocking was performed at room temperature for 1 hour using a M.O.M. blocking solution (#BMK-2202; Vector Laboratories, Burlingame, CA, USA) supplemented with 5% bovine serum albumin (BSA) in 1× PBS to reduce nonspecific antibody binding. Primary antibody staining was performed by incubating sections overnight at 4°C with antibodies diluted in M.O.M. diluent with 5% BSA. The following primary antibodies were used: anti-Ror2 (1:500; Developmental Studies Hybridoma Bank, Iowa City, IA), rabbit anti-eGFP (1:250; #ab290; Abcam, Cambridge, MA, USA), mouse anti-eGFP (1:250; JL-8, #632381; Takara, Kusatsu, Shiga, Japan), anti-RFP (1:250; #600-401-379; Rockland, Pottstown, PA, USA), anti-K8 (1:250; #TROMA-1; Developmental Studies Hybridoma Bank), anti-K14 (1:5,000; #905301; BioLegend, San Diego, CA, USA), anti-K5 (1:5,000; #905501; BioLegend), anti-p63 (1:200; ΔN-p63, #619002; BioLegend), anti-BrdU (1:250; #ab6326; Abcam), anti-ERα (1:250; #sc-542; Santa Cruz Biotechnology, Dallas, TX, USA), anti-PR (1:500; #A0321; ABclonal Technology, Woburn, MA, USA), and anti-YAP1 (1:500; #14074; Cell Signaling Technology, Danvers, MA, USA). After three PBS washes, sections were incubated with secondary antibodies for 1 hour at room temperature in the dark. Alexa Fluor 488–, 594–, or 647–conjugated goat anti-rabbit, anti-mouse, anti-rat, or anti-chicken IgG secondary antibodies (ThermoFisher Scientific) were diluted in M.O.M. diluent containing 5% BSA. Sections were washed three times with 1× PBS and counterstained with 1 μg/ml 4′,6-diamidino-2-phenylindole (DAPI) before mounting with ProLong Diamond Antifade Mounting Media (#P36961; ThermoFisher Scientific). For Ror2 detection, tyramide signal amplification was performed according to the manufacturer’s instructions (#NEL701A001KT; PerkinElmer, Waltham, MA, USA).

### Western Blotting

Proteins were separated on NuPAGE 4–12% or 10% Bis-Tris protein gels (ThermoFisher Scientific, #NP0336BOX and #NP0315BOX) using MOPS running buffer, following the manufacturer’s protocol. The separated proteins were transferred onto polyvinylidene fluoride (PVDF) membranes (ThermoFisher Scientific, #LC2002) using a wet transfer system to ensure efficient protein transfer. Membranes were subsequently blocked at room temperature for 1 hour in 5% blocking reagent (Bio-Rad, #1706406; Hercules, CA) dissolved in Tris-buffered saline containing 0.05% Tween-20 (TBST) to minimize nonspecific antibody binding. Blocked membranes were incubated overnight at 4°C with primary antibodies diluted in TBST under gentle agitation. After primary antibody incubation, the membranes were washed three times with TBST for 5 minutes per wash and then incubated with horseradish peroxidase (HRP)–conjugated secondary antibodies diluted in TBST for 1 hour at room temperature. Following secondary antibody incubation, membranes were washed three additional times in TBST. Protein bands were visualized using an enhanced chemiluminescence (ECL) substrate, and the signals were detected using a chemiluminescence imaging system. The following primary antibodies were used, with their respective dilutions indicated: Ror2 (1:1000; Developmental Studies Hybridoma Bank, Iowa City, IA), RhoA (1:1000; #2117; Cell Signaling Technology [CST], Danvers, MA), ROCK1 (1:1000; #4035; CST), Cdc42 (1:1000; #2466; CST), Rac1/2/3 (1:1000; #2465; CST), YAP/TAZ (1:1000; #8418; CST), phosphorylated YAP (p-YAP, 1:1000; #13008; CST), LATS1 (1:1000; #3477; CST), phosphorylated LATS1 (p-LATS1, 1:1000; #9157; CST), and GAPDH (1:2500; #5174; CST). HRP-conjugated secondary antibodies specific to rabbit or mouse were sourced from Bio-Rad.

### RNA Isolation, Sequencing, and Analysis

Total RNA was extracted and purified from samples using the RNeasy Mini Kit (Qiagen, #74104; Germantown, MD) according to the manufacturer’s instructions. RNA quality and concentration were initially assessed using the Nanodrop ND-1000 spectrophotometer (ThermoFisher Scientific, Waltham, MA) and further evaluated for integrity with an Agilent Bioanalyzer Nano chip (Agilent Technologies, Santa Clara, CA). These quality checks were performed by the Baylor College of Medicine (BCM) Genomic and RNA Profiling Core to ensure RNA samples met the criteria for downstream applications. RNA integrity numbers (RINs) of ≥7.0 were required for sequencing. RNA-Seq library preparation, including poly(A) enrichment and cDNA synthesis, was conducted by the BCM Genomics and RNA Profiling Core. Libraries were sequenced on the Illumina NovaSeq 6000 platform, producing paired-end reads of 100 base pairs to achieve a target depth of at least 40 million reads per sample. Post-sequencing, read alignment to the reference genome was performed using STAR or HISAT2 algorithms, ensuring high alignment accuracy. Transcript abundance was quantified using Cufflinks, and expression values were calculated as fragments per kilobase of transcript per million mapped reads (FPKM). Prior to downstream analysis, FPKM values were log₂-transformed. Differential expression analysis was conducted to identify gene expression changes between wild-type (WT) and Ror2 knockout (KO) groups. Genes were considered significantly altered under KO conditions based on an adjusted p-value of <0.05 and a fold-change threshold of >2 (upregulated) or <0.5 (downregulated). The Database for Annotation, Visualization and Integrated Discovery (DAVID) was used to help to prioritize gene sets by identifying enriched biological themes, functional-related gene groups, and interacting proteins that were differentially expressed^53^. These analyses highlighted enriched biological processes, functional gene groups, and interacting proteins. Results were used to prioritize gene sets for further investigation of biological themes relevant to the experimental context. RNA sequencing data have been deposited in NCBI’s Gene Expression Omnibus (GEO) repository under accession numbers [Pending].

### Quantitative Real-Time PCR

Total RNA was reverse-transcribed into complementary DNA (cDNA) using the High-Capacity RNA-to-cDNA Kit (Applied Biosystems, #4388950; Foster City, CA) following the manufacturer’s protocol. Each reaction was performed with 1 µg of total RNA in a final reaction volume of 20 µl. The resulting cDNA was diluted to 10 ng/µl before use in quantitative PCR reactions. Quantitative real-time PCR (qPCR) was carried out using the StepOnePlus Real-Time PCR System (Applied Biosystems) with SYBR Green PCR Master Mix (GenDEPOT, #Q5602-050; Baker, TX) as the detection reagent. Primer sequences (detailed in Supplemental Table S1) were custom-designed and purchased from Sigma-Aldrich (KiCqStart SYBR Green Primers). Each reaction was performed in triplicate in a 25 µl volume, including 5 µl of cDNA, 12.5 µl of SYBR Green Master Mix, and 500 nM of each primer.

### FACS Staining and Analysis

Lineage-positive cells were depleted from single-cell preparations using the EasySep Mouse Mammary Stem Cell Enrichment Kit (StemCell Technologies, #19757) following the manufacturer’s protocol. The remaining cells were resuspended at a concentration of 10⁷ cells/ml in HBSS^+^ (Hank’s Balanced Salt Solution supplemented with 2% fetal bovine serum [FBS] and 10 mM HEPES buffer) to maintain viability during antibody staining.

Antibody staining was conducted on ice for 30 minutes to minimize nonspecific binding and maintain cellular integrity. Cells were labeled with anti-mouse CD24-Pacific Blue (1:100; BioLegend, #101819) and anti-mouse CD29-APC (1:100; BioLegend, #102215) to distinguish luminal and basal epithelial populations. Separation of myoepithelial (basal) and luminal fractions was based on CD24 and CD29 expression levels. Following staining, cells were washed twice with HBSS^+^ and filtered through a 40 µm strainer to ensure single-cell suspensions for downstream analysis. Flow cytometry was performed using either an LSR Fortessa cell analyzer (BD Biosciences) or an Aria II cell sorter (BD Biosciences) for analysis or sorting, respectively. Compensation controls were included to correct for spectral overlap between fluorophores, and data acquisition was performed with BD FACSDiva software. Subsequent data analysis was conducted using FlowJo software, version 10.9.0. All fluorescence-activated cell sorting (FACS) plots depict compensated data that have undergone display transformation.

### 10x Genomics Single-Cell ATAC-Sequencing and Data Analysis

Lineage-depleted, FACS-sorted GFP-positive cells were used for single-cell ATAC-sequencing. Cells were washed and resuspended in PBS containing 0.04% BSA (PBS^+^). Cell quantity and viability were assessed using Acridine Orange and Propidium Iodide dyes on a Cellometer Auto 2000 (Nexcelom). Samples with a viability of >50% and a total of 100,000–1,000,000 cells underwent nuclei isolation using the 10x Genomics Nuclei Isolation for Single-Cell ATAC-Sequencing Kit, following the manufacturer’s protocol. Isolated nuclei were quantified, and approximately 15,000 nuclei were loaded onto a Chromium X (10x Genomics) platform using the Chromium Next GEM Single Cell ATAC Reagent Kit v2. Prior to nuclei partitioning and GEM generation, nuclei suspensions were transposed. The GEM contents were then pooled and purified using magnetic beads. DNA libraries were amplified for seven cycles and further purified using SPRI beads (Beckman Coulter). Library quality was assessed using a High Sensitivity D1000 Tapestation (Agilent Technologies) and quantified using the Qubit 2.0 DNA HS assay (ThermoFisher Scientific). Equimolar pooling of libraries was performed based on QC metrics, and the samples were sequenced on an Illumina® NovaSeq X Plus platform with a paired-end read length configuration of 150 bp, targeting 1 billion paired-end reads per sample (0.5 billion reads in each direction).

Raw sequencing data were analyzed using Cell Ranger ATAC pipelines (version 2.1.0). FASTQ files were processed using the cellranger-atac count function, which included read alignment, cell barcode counting, and identification of transposase cut sites. After initial processing, the cellranger-atac aggr function was used to aggregate data from the p63-WT and p63-KO groups. Aggregation analysis included normalization of sequencing depth to achieve the same median fragments per cell across samples, identification of accessible chromatin peaks, and generation of count matrices for chromatin peaks and transcription factor binding sites.

Processed data underwent dimensionality reduction using t-SNE or UMAP to visualize cell clusters based on chromatin accessibility profiles. Downstream analyses included differential chromatin accessibility analysis, transcription factor motif analysis, and promoter accessibility profiling to identify differences between experimental groups. Visualization of results was performed using the 10x Genomics Loupe Browser (version 7). Additional data visualizations, including heatmaps and volcano plots, were generated in R (version 4.3.3) using the ggplot2 package.

### Lentiviral Ror2 Knockdown and Wnt5a Treatment in Basal Mammary Epithelial Cells In Vitro

Lentiviral vectors were generated using LeGO plasmids encoding short hairpin RNA (shRNA) targeting either luciferase (shLUC, control) or Ror2 (shRor2). The shRNA antisense sequences were as follows: shLUC, 5′-ATTCCAATTCAGCGGGGGC-3′; and shRor2, 5′-TATTCTGCGTAAAGCACCACG-3′. The shRor2 sequence corresponds to the shRor2-94 clone from the MISSION pLKO Lentiviral shRNA library (Sigma-Aldrich, St. Louis, MO). All constructs were sequence-validated to confirm accuracy. Primary basal mammary epithelial cells were isolated and FACS-sorted as described previously. Approximately 500,000 basal cells per well were infected overnight (16 hours) in suspension using ultra-low attachment 24-well plates at a multiplicity of infection (MOI) of 30. Following infection, cells were washed three times with PBS to remove residual virus and seeded into 6-well tissue culture plates for further treatment. Recombinant human/mouse Wnt5a protein (100 ng/ml; Bio-Techne, #645-WN; Minneapolis, MN) was added to half of the wells containing shLUC or shRor2 cells 72 hours post-transduction. Treated cells were incubated with Wnt5a for 16 hours before collection. Cell lysates were harvested using RIPA buffer containing protease and phosphatase inhibitors for subsequent analyses, including quantitative real-time PCR, Western blotting, and RhoA activation assays.

### RhoA Activation Assay

The RhoA activation pull-down assay was conducted using the RhoA Activation Assay Biochem Kit (Cytoskeleton, Inc., #BK036-S; Denver, CO) according to the manufacturer’s protocol. Protein lysates were prepared from shLUC and shRor2 lentiviral-transduced basal mammary epithelial cells, with or without Wnt5a treatment, using the lysis buffer provided in the kit supplemented with protease and phosphatase inhibitors.

Cell lysates were clarified by centrifugation at 12,000 × g for 10 minutes at 4°C. Equal amounts of protein lysate, as determined by the Bradford Protein Assay (Bio-Rad, #5000006; Hercules, CA), were incubated with Rhotekin-RBD (Rho-binding domain) protein-conjugated agarose beads to selectively pull down active RhoA-GTP. Incubations were performed at 4°C with gentle rotation for 1 hour. After incubation, beads were washed three times with the wash buffer provided in the kit to remove non-specifically bound proteins. The bound RhoA-GTP was eluted from the beads and analyzed by SDS-PAGE, followed by Western blotting. Detection was performed using a RhoA-specific primary antibody (1:1000; Cell Signaling Technology, #2117) and an HRP-conjugated secondary antibody. GAPDH expression in total protein lysates was used as a loading control to confirm equal protein input across samples. Western blot signals were visualized using an enhanced chemiluminescence (ECL) substrate, and band intensities were quantified using ImageJ software.

### Statistical Analysis and Rigor

All data are presented as the mean ± standard deviation (SD). The number of biological replicates (n) is specified in the figure legends unless otherwise indicated. For comparisons involving more than two groups, statistical significance was determined using one-way analysis of variance (ANOVA) followed by Tukey’s multiple comparison test to account for multiple comparisons. For two-group comparisons, unpaired two-tailed Student’s t-tests were performed unless stated otherwise. Statistical analyses were conducted using GraphPad Prism 10 or FIJI (ImageJ). Significance thresholds were set at p < 0.05 for all tests, with levels of significance denoted as follows: p < 0.05 (*), p < 0.01 (**), p < 0.001 (***), and p < 0.0001 (****). To ensure rigor and reproducibility, all experiments were independently reproduced in at least three biological replicates, with consistent results observed across replicates. Where applicable, technical replicates were also included to further validate findings. Variations in sample sizes or statistical tests applied are detailed in the respective figure legends.

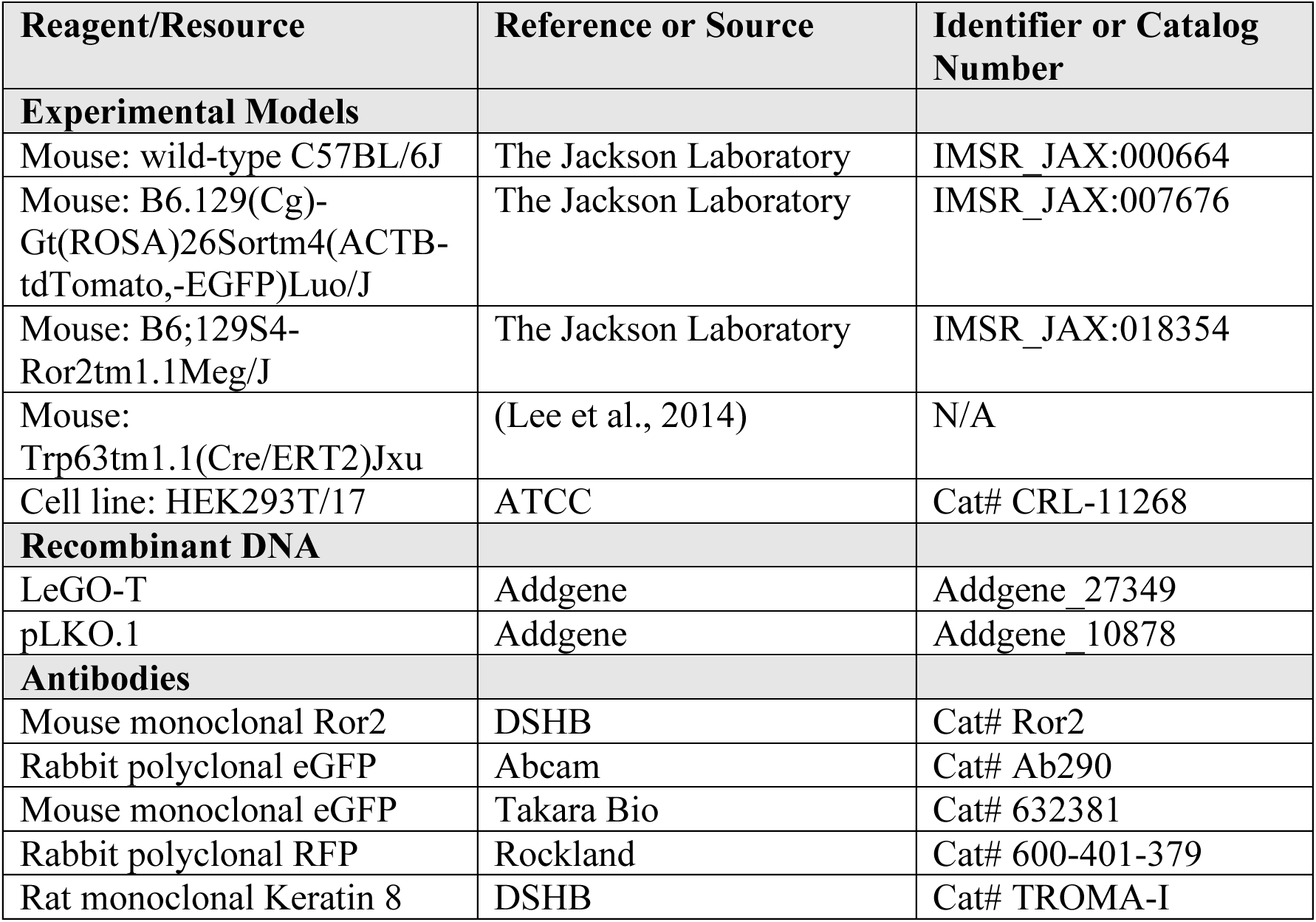
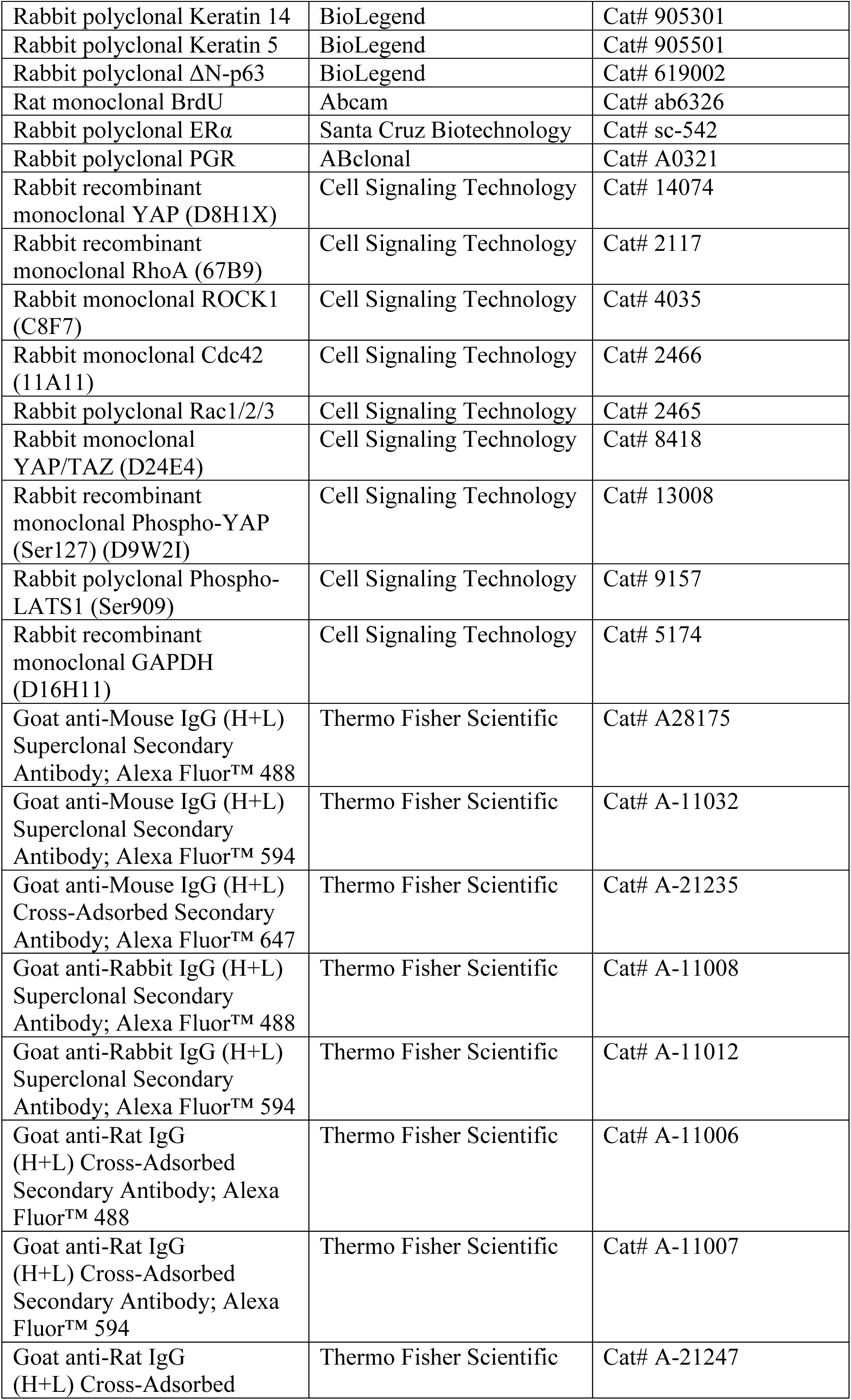
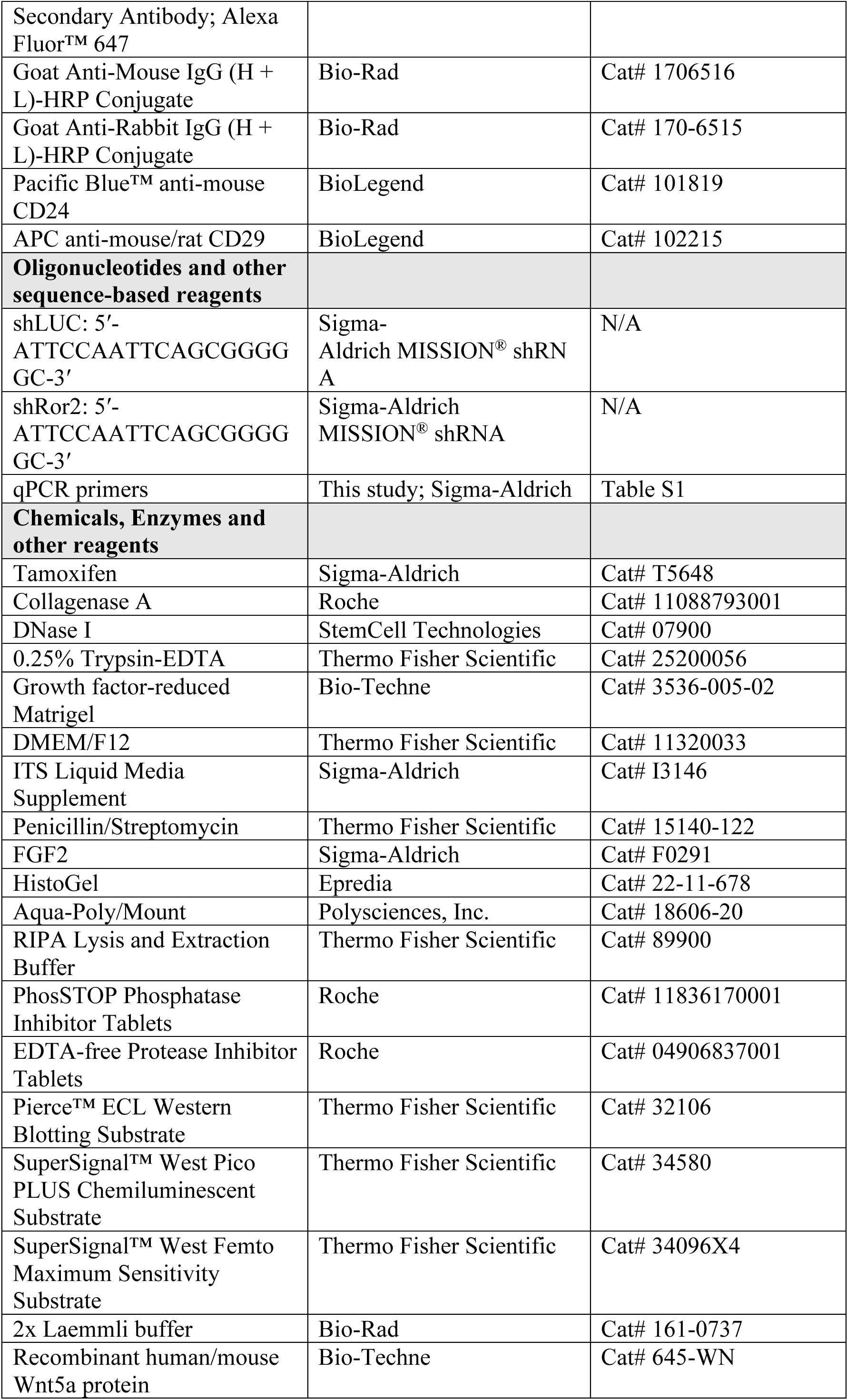
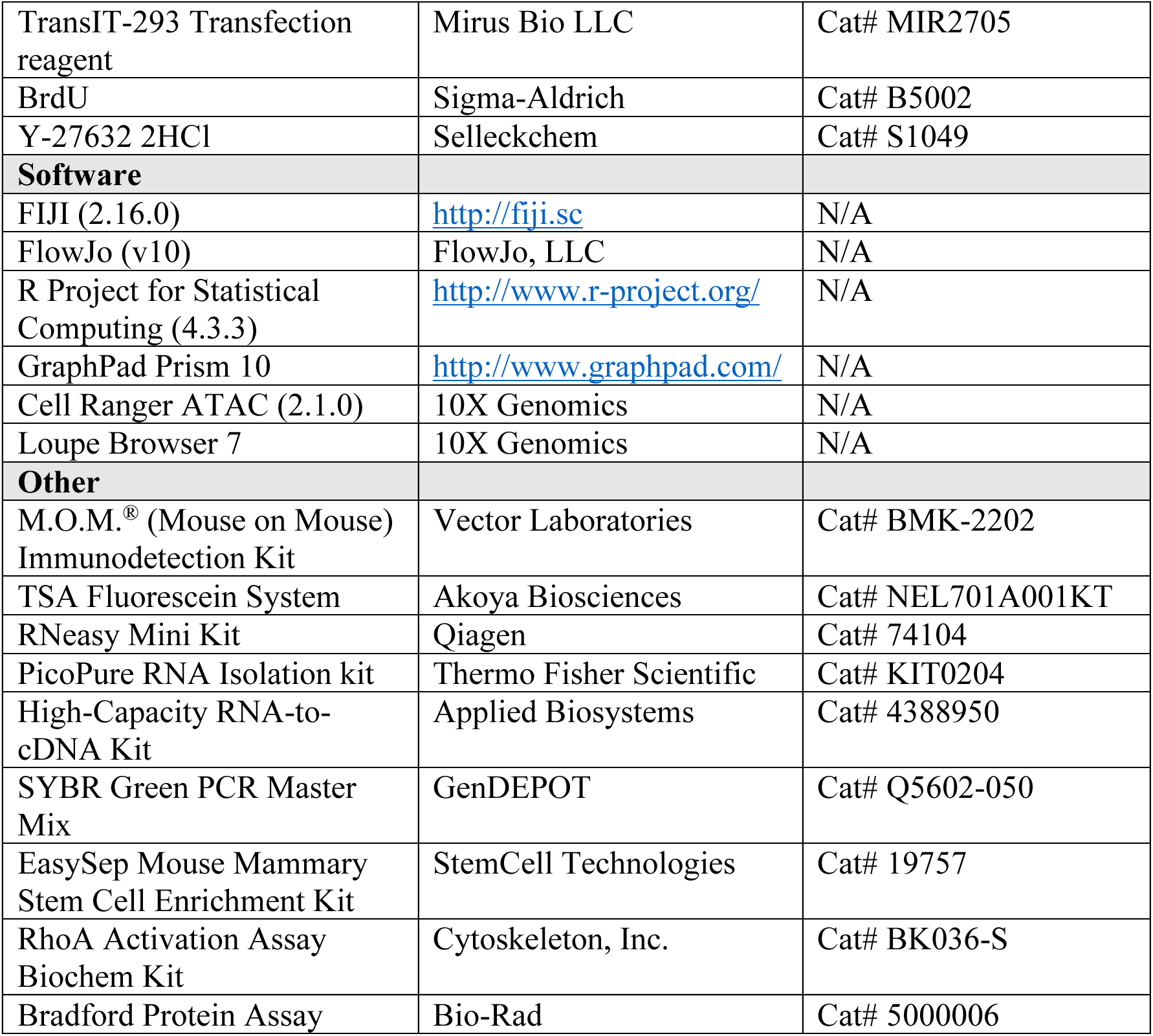
Reagents and Tools Table.

## Acknowledgements

We extend our gratitude to Jeffrey M. Rosen of Baylor College of Medicine for his invaluable insights and constructive discussions. This work was supported by 5R01ES033601 from the National Institute of Environmental Health Sciences, 5K22CA207463 from the National Cancer Institute (NCI), Susan G. Komen CCR18548284, and the L. E Gordon Cancer Research Fund, the Caroline Wiess Law Fund for Research in Molecular Medicine to K.R. H. S. was supported by RP210027 from the Cancer Prevention and Research Institute of Texas (CPRIT) and CA016303-46. We thank Fengju Chen for bioinformatics support within the Biostatistics and Cancer Bioinformatics Division of the Dan L. Duncan Comprehensive Cancer Center. This project was also supported by the Genomic and RNA Profiling Core at the Baylor College of Medicine and the Cytometry and Cell Sorting Core at the Baylor College of Medicine with funding from Cancer Prevention and Research Institute of Texas (CPRIT) Core Facility Support Award CPRIT-RP180672 and National Institutes of Health (NIH) grants P30 CA125123 and S10 RR024574. Imaging for this project was supported by the Optical Imaging & Vital Microscopy (OiVM) core at Baylor College of Medicine.

## Author Contributions

H.S. and K.R. wrote the manuscript. K.R. edited the manuscript, conceived the study, and analyzed the data in collaboration with H.S. H.S. carried out all experiments and analyzed the data. E.M.M. and M.E. contributed to immunostaining, imaging, and mouse experiments. C.C. performed the analysis of bulk RNA sequencing data. J.X. developed the p63^CreERT2^ knock-in mouse.

