## Supplementary figures and images for "Noncanonical Wnt/Ror2 Signaling Regulates Basal Cell Fidelity and Branching Morphogenesis in the Mammary Gland"

### Supplemental Figure 1

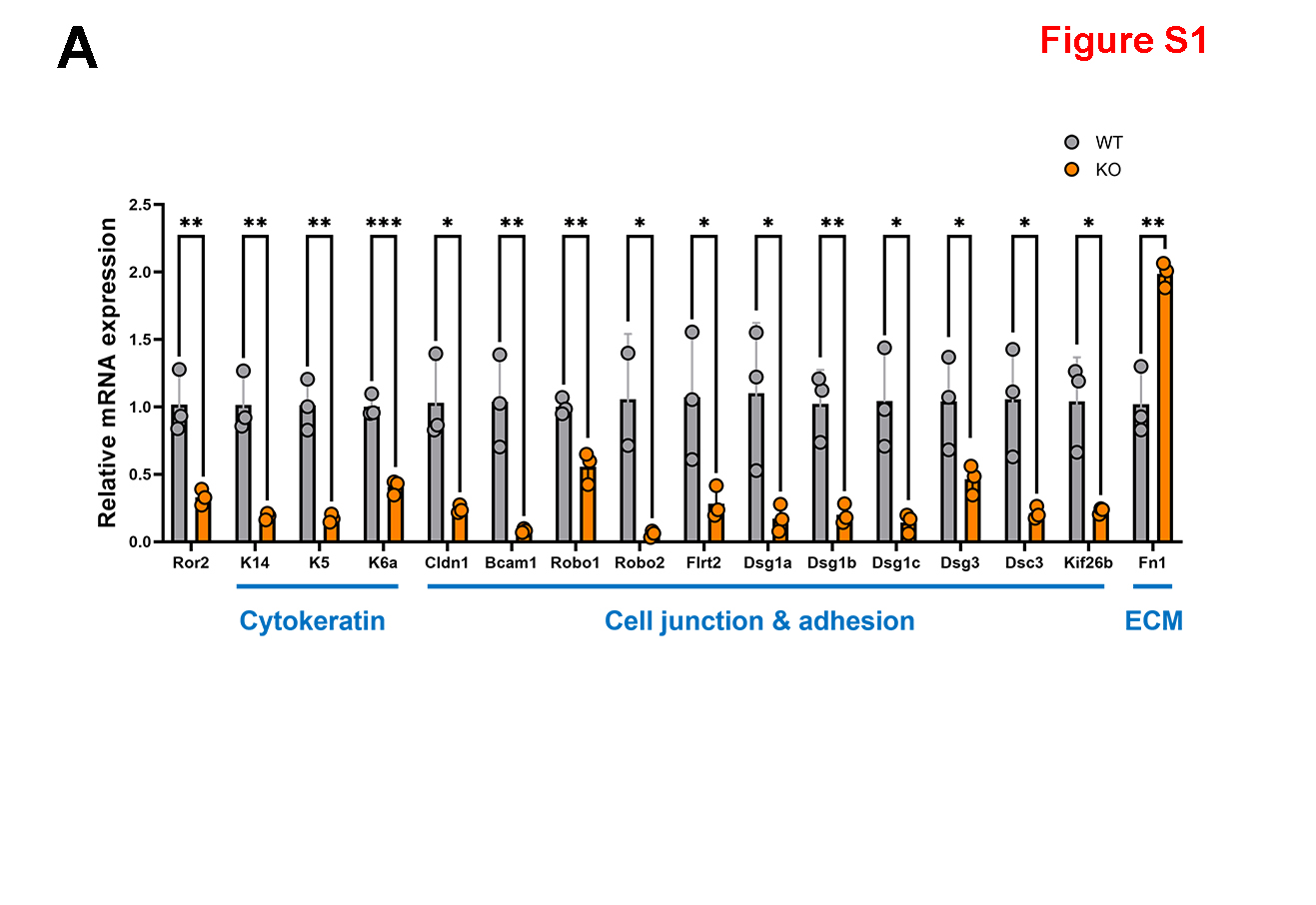

### Supplemental Figure 2

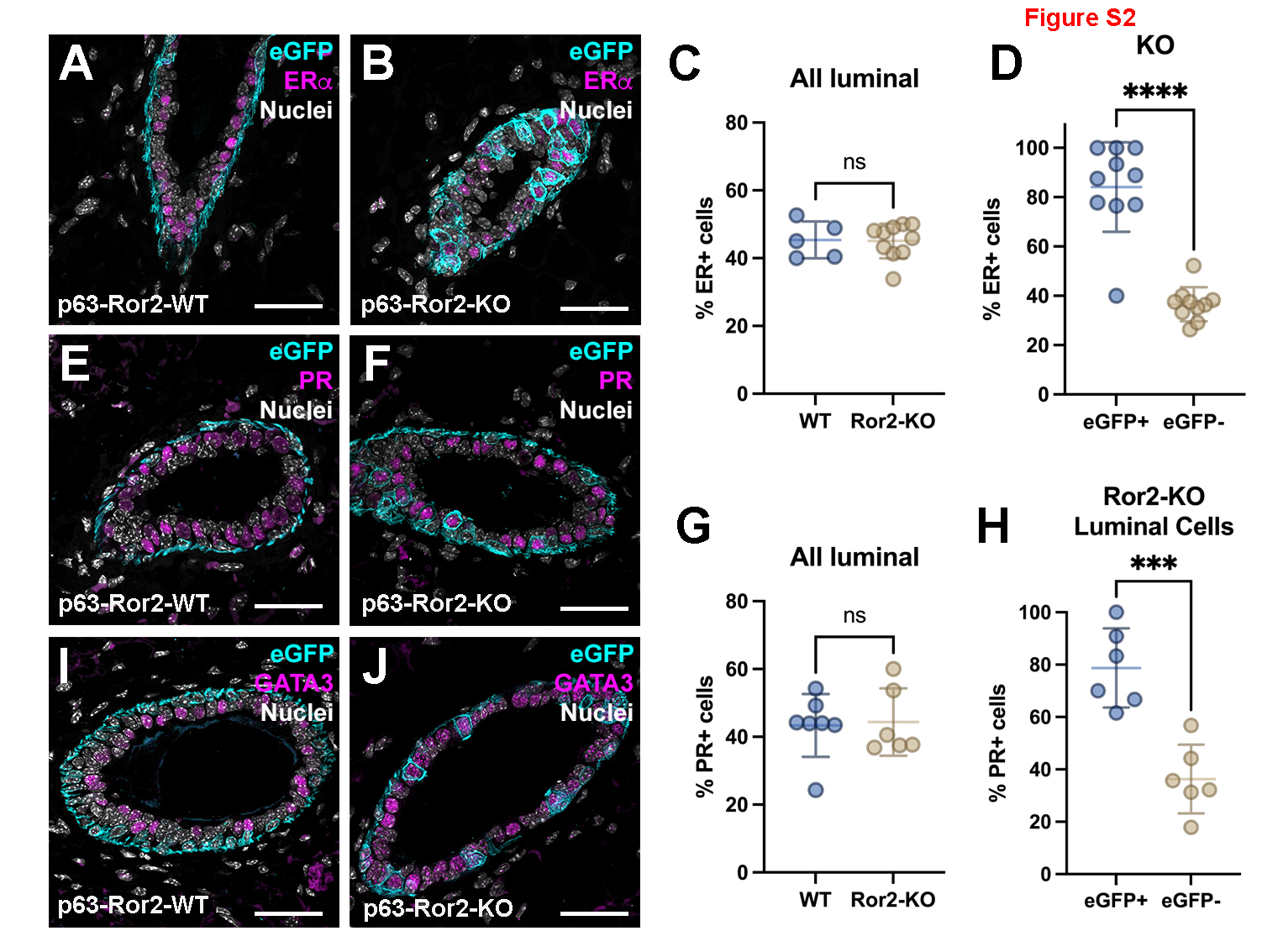
